# Rab5 Overcomes CAR T Cell Dysfunction Induced by Tumor-Mediated CAR Capture

**DOI:** 10.1101/2024.07.26.605334

**Authors:** Meidi Gu, Edmund J. Carvalho, Kaitlin A. Read, David Padron Nardo, James L. Riley

**Author notes:** **Corresponding authors:** Meidi Gu James L. Riley.

## Abstract

Continuous interaction between chimeric antigen receptor (CAR) T cell (CART) and tumors often result in CART dysfunction and tumor escape. We observed that tumors can take up CAR molecules, leaving CARTs without surface-expressed CARs and thus unable to kill tumors after prolonged exposure. Overexpression of Rab5 resulted in augmented clathrin-independent endocytosis, preventing loss of surface-expressed CARs, and enhanced CART activity. Interestingly, we observed membrane protrusions on the CART cell surface which disappeared after multiple tumor challenges. Rab5 maintained these protrusions after repeated tumor engagements and their presence correlated with effective tumor clearance, suggesting a link between endocytosis, membrane protrusions, and cytolytic activity. *In vivo*, Rab5-expressing CARTs demonstrated improved activity and were able to clear an otherwise refractory mesothelin-expressing solid cancer in humanized mice by maintaining CAR surface expression within the tumor. Thus, pairing Rab5 with CAR expression could improve the clinical efficacy of CART therapy.

**Highlights**

- “CAR-jacking” occurs when surface CAR is internalized by target tumor cells.
- Rab5 overexpression prevents “CAR-jacking” and enhances CART function.
- Rab5 promotes CAR endocytic recycling and maintains membrane protrusions.
- Rab5-expressing CARTs exhibit enhanced therapeutic efficacy against solid tumors.

## INTRODUCTION

CART therapy has led to remarkable therapeutic responses in patients with certain hematological malignancies^1^. However, to date, CARTs have not achieved the same impact on non-hematological tumors^2,3^. Significant effort has been devoted to boosting the efficacy of CARTs to expand the number of patients who could benefit from this therapy, including providing signals that prolong CART persistence^4–7^, resist T cell exhaustion^8^, and help CARTs overcome the immunosuppressive tumor microenvironment^9^. CARs leverage the modular aspect of immune receptors, enabling the construction of synthetic receptors that integrate the functional properties of their component parts^10^. While most of these properties promote the ability of the CAR to redirect the immune system, some have been found to limit CART function. For example, trogocytosis is a process whereby T cells capture and internalize peptide-major histocompatibility complex protein complexes (pMHCs) through their TCRs^11^. CARs follow a similar fate as TCRs, undergoing trogocytosis after antigen recognition, and this uptake of tumor antigen makes the CART susceptible to fratricide and thus promotes tumor escape^12^. Using low-affinity CARs or CTLA4-cytoplasmic tail-fused CARs has been shown to minimize CAR-mediated trogocytosis and enhance CART antitumor efficacy^13,14^. Additionally, following CAR internalization into CARTs, Li and colleagues demonstrated that CARs signaling domains undergo ubiquitination, leading to their lysosomal degradation and downregulation. As such, blocking this process enhances CAR expression and CART activity^15^. Both examples illustrate that CAR trafficking after target engagement plays a role in limiting CART activity. Another mechanism by which immunoreceptors are cleared from the cell membrane is ectocytosis, where TCRs are captured by target cells, preventing excessive activation of TCR cascade^16^. In this manuscript, we found that profound CAR loss occurs in a similar manner following CART interaction with tumor cells, a process we have termed ‘‘CAR-jacking’’. It joins T cell exhaustion^17^, target loss^18^ and the tumor microenvironment^19^ as a major cause of CART dysfunction.

Rab GTPases are key organizers of membrane trafficking^20^. They are commonly used as markers and identifiers of various cargos in the endocytic and secretory systems^21^. The human genome encodes more than 60 Rab GTPases. Among them, Rab5, an early endosome marker, plays a crucial role in regulating the internalization and trafficking of membrane receptors including TCRs^22–24^. Mutations in the Rab5 gene directly affect TCR transport and result in aberrant, albeit enhanced TCR signaling^25–27^. Given that TCR activation results in local and temporary inhibition of endocytosis, which is required for ectocytosis to occur^16^, we hypothesized that enhancing endocytic activity may prevent ectocytosis and ultimately CAR loss. To test this hypothesis, we overexpressed Rab5 and demonstrated that it effectively interfered with “CAR-jacking”. Rab5 facilitated ’cis-internalization’ of CARs by CARTs, which greatly limited tumor cell ’trans-internalization’-mediated CAR loss. Collectively, our data show that “CAR-jacking” can result in significant CART dysfunction, and that this can be overcome by overexpressing Rab5, providing both the basis and rationale to explore Rab5 overexpression in CARTs in the clinic.

## RESULTS

### Surface CAR expression is lost after repetitive tumor challenge

To better understand how tumor cell and CART interactions alter long term CART function, we employed a continuous antigen exposure (CAE) assay where tumor cells are periodically mixed with CARTs to better mimic *in vivo* conditions^28^. Primary human T cells were activated with CD3/28 coated beads, transduced with a CD19-specific CAR that employed the 4-1BB and CD3 zeta signaling domains (19BBz), and cultured for 9 days. CARTs were then repetitively challenged with K562 expressing huCD19 and GFP (K.19.GFP) every two days. During this time, CAR expression and tumor killing were monitored (Figure S1A). We observed that CAR expression on both CD4^+^ and CD8^+^ T cells significantly diminished after the initial challenge (Figure 1A), regardless of the detection method (Figure S1B). Although CAR expression initially resurfaced as previously described^15,28^ (Figure 1A), it failed to reappear after prolonged stimulation (3^rd^-5^th^ challenge) (Figure 1A). The inability of CARTs to maintain CAR expression correlated with a reduced tumor cell killing capacity (Figure 1B and Figure S1C). Similar results were obtained using a CAR that employed the CD28 costimulatory domain or a mesothelin-specific CAR (Figure S1D, S1E), indicating that this phenomenon is general property of CARTs. These results reinforce previous research showing that loss of CAR surface expression significantly limits CART function^29,15,30^ and extend these studies by demonstrating that CARTs lose their ability to maintain surface CAR expression after extended tumor exposure.

**Figure 1.**
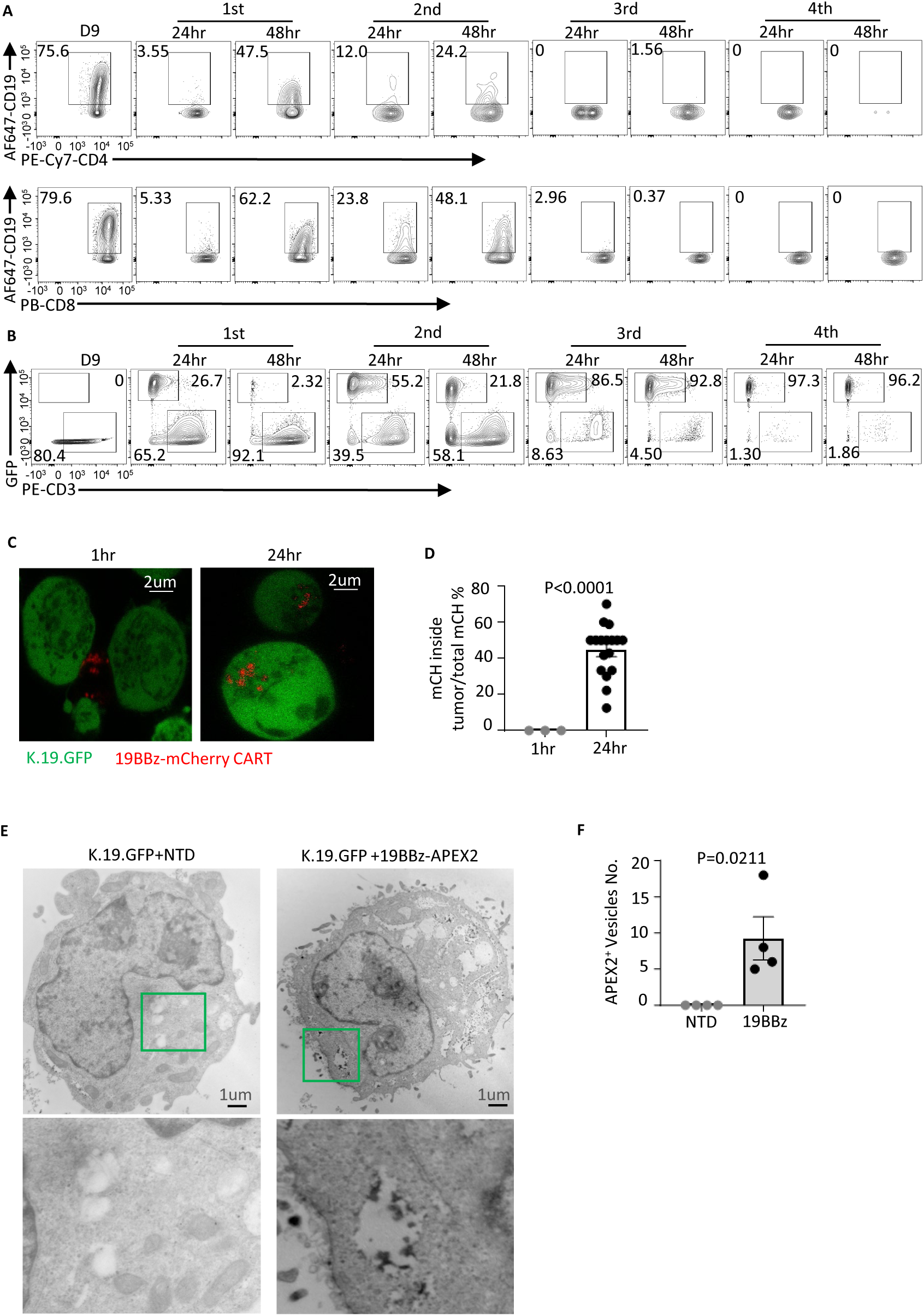
Surface CAR expression is lost after repetitive tumor challenge. (A). Primary human T cells were activated and transduced with a 19BBz CAR and cultured for 9 days. CAE assay was performed by co-culturing 1x10^6^ CAR positive T cells with 2 x10^6^ K562 cells expressing CD19 and GFP (K.19.GFP). Every 2 days, additional fresh 2x10^6^ K.19.GFP cells were added to the co-culture assay and surface CAR expression on CD4 and CD8 T cells was measured by flow cytometry. (B). The presence of tumor (GFP^+^) and T cells (CD3^+^) in the CAE assay was measured by flow cytometry on the indicated time points. (C). Confocal imaging of the cellular localization of CAR. CARTs with mCherry (mCH) fused to the CD3ζ chain incubated with K.19.GFP cells at 1:1 ratio as the indicated times prior to live cell imaging. (D). Summary data showing percentage of mCherry-CAR internalized into tumor cells as shown in (C). (E). Transmission electron microscopy (TEM) imaging of tumor cells capturing APEX2 fused 19BBz-CAR (black) to the intracellular vesicles. Images were captured on K.19.GFP cells 24-hours after the third-round re-challenge assay. T cells were depleted by CD3 beads prior to DAB staining. (F). Quantification of APEX2-CAR containing vesicles inside of K.19.GFP tumor cells as shown in (E). Experiments shown in (A)-(D) are representative of three independent experiments, in (E) and (F) are representative of two independent experiments. Error bar show mean ± SEM. P value was determined by Students t test. See also Figure S1.

To further investigate the role of tumor plays in the loss of CAR expression, CARTs were co-cultured with antigen-expressing tumor cells at different ratios. We found that an increased ratio of tumor cells:CARTs led to greater loss of surface CAR (Figure S1F, S1G), and that was dependent upon direct tumor-CARTs interactions (Figure S1H). To explore whether CARs were captured by tumor cells, similar as observed when TCRs interact with target cells expressing cognate pMHC^16^, we generated functional CARTs with mCherry fused to the C-terminus of the CD3 zeta chain and confirmed that these mCherry-fused CARTs functioned similarly to those lacking mCherry (Figure S1I). Co-culturing K.19.GFP tumor cells with mCherry-CARTs revealed that tumor cells captured CARs from the T cells after 24 hours incubation (Figure 1C, 1D). To further understand the fate of CAR after CARTs are co-cultured with tumors, we attached pea ascorbate peroxidase 2 (APEX2) to the CAR construct and imaged CAR location with transmission electron microscopy (TEM). We observed APEX2-CAR transferring from T cells, crossing cell junctions (Figure S1J), and being internalized into the vesicles within tumor cells (Figure 1E, 1F). This process, which we have termed ‘CAR-jacking’, represents a novel form of CART cell dysfunction caused by tumor-mediated CAR internalization. Ultimately, this leads to CARTs being unable to maintain surface CAR expression and function after extended tumor interactions.

### Rab5 maintains surface CAR expression and CAR T cell function

As previously described, endocytosis of an activated TCR is inhibited, which subsequently allows ectocytosis across the synapse^16^. We postulated that enhancing the endocytic activity of CARTs could allow them to retain CAR molecules and prevent their transfer into target tumor cells. Rab5 is the key component of early endosomes and plays a crucial role in endocytosis^24^. We initially examined Rab5 expression in T cells and found it was not detected in resting T cells. However, we observed transient upregulation of Rab5 following T cell activation, which diminished as the T cells were cultured (Figure 2A). We hypothesized that sustained expression of Rab5 would boost endocytic activity, allowing CARTs to retain CARs during continuous tumor challenge. To test this, we generated 19BBz CAR vectors co-expressing either truncated EGFR (19BBz-tEGFR) as a control or Rab5 (19BBz-Rab5). We observed that 19BBz-Rab5 CARTs maintained high Rab5 expression throughout the culture period, whereas Rab5 expression was undetectable in 19BBz-tEGFR CARTs (Figure 2B). We then compared 19BBz-Rab5 and 19BBz-tEGFR CARTs in the CAE assay. Remarkably, CAR expression on 19BBz-Rab5 CARTs persisted after five tumor challenges, whereas 19BBz-tEGFR CART lost CAR surface expression after three challenges in both CD4^+^ T cells (Figure 2C,2D) and CD8^+^ T cells (Figure S2A, S2B). This sustained CAR expression led to efficient tumor clearance, demonstrating that persistent Rab5 expression greatly extends CARTs anti-tumor function (Figure 2E,2F). EMMESO, a human mesothelioma cell line isolated from a patient’s tumor and stably transduced to express human mesothelin, is a challenging tumor to control both *in vitro* and *in vivo*^31,32^ . We therefore tested whether Rab5 enhances CART activity against EMMESO in CAE assay (Figure S2C). Mesothelin-specific Rab5 CARTs maintained robust CAR expression (Figure 2G,2H). However, unlike CD19 expressing tumors, EMMESO cells were never completely cleared from the culture by MESOBBz-tEGFR CARTs lacking Rab5. Impressively, MESOBBz-Rab5 CARTs successfully cleared EMMESO cultures after seven challenges (Figure 2I,2J), indicating that Rab5 expression may be especially critical for CARTs targeting solid tumors. Collectively, these findings demonstrate that Rab5 overexpression promotes durable CAR surface expression and CART antitumor activity.

**Figure 2.**
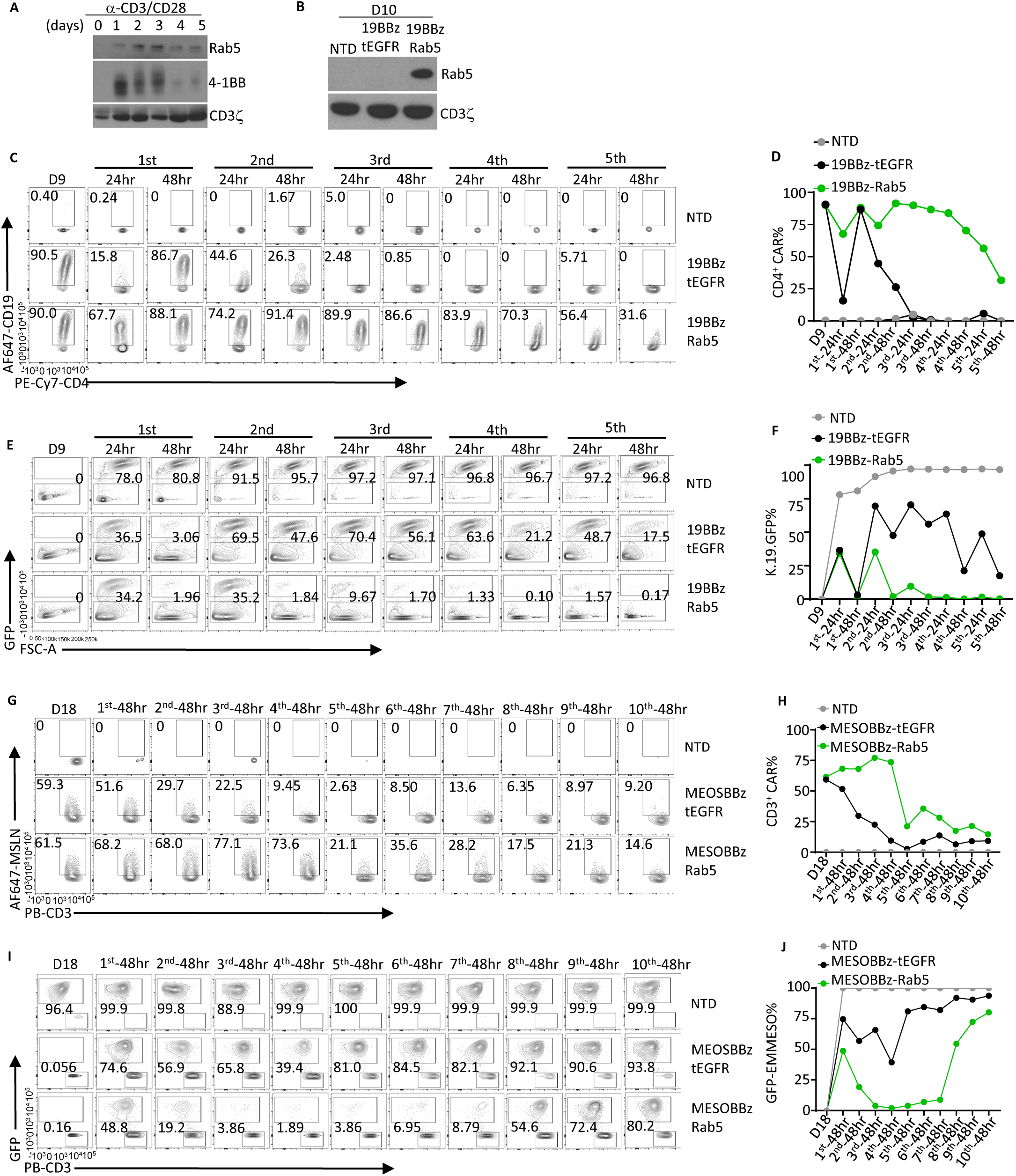
Rab5 enhances surface CAR persistence and CART function. (A). Human primary T cells were stimulated with CD3/CD28 coated beads for three days, whole cell lysates were collected on indicated days. Rab5, 4-1BB (T cell activation control) and CD3 zeta chain (loading control) were detected by immunoblotting. (B). Activated primary human T cells were not transduced (NTD) or transduced with a lentiviral vector expressing 19BBz-CAR linked to tEGFR (19BBz T2A tEGFR) or one that co-expressed Rab5 (19BBz T2A Rab5). After 10 days of culture, whole cell lysates were collected and immunoblotting for Rab5 and CD3 zeta was performed. (C-F). CAE assay shown in Figure S1A was performed by co-culturing K.19.GFP with 19BBz-tEGFR or 19BBz-Rab5 CARTs. The ability to bind CD19 protein in CD4^+^ CARTs (C, D) and presence of GFP expressing tumors (E, F) at the indicated times was determined by flow cytometry. (G-J). Non-transduced (NTD), mesothelin-specific CARTs (MESOBBz-tEGFR), Rab5 expressing mesothelin-specific CARTs (MESOBBz-Rab5) were challenged in the CAE assay shown in Figure S2C with GFP-transduced EMMESO cells. Labeled mesothelin protein was used to test the MESO-CAR expression at the indicated time points (G, H). The presence of GFP^+^ EMMESO cells was measured the indicated time points (I, J) by flow cytometry. Data is representative of three independent experiments. See also Figure S2.

#### Rab5 requires a properly regulated GTPase to prevent CAR loss

To determine whether Rab5 provides survival signals that allow CARTs to preferentially expand during the CAE assay, we transduced T cells with CARs with mCherry placed after the CD3 zeta chain in the presence and absence of Rab5 and placed these T cells in the CAE assay. We again observed marked differences in CAR surface expression (Figure 3A, 3C), while the frequency of T cells expressing mCherry (CARTs) remained similar (Figure 3B,3C), suggesting that Rab5 expression does not preferentially expand or enrich for CARTs but rather allows CARTs to maintain CAR surface expression. We also evaluated whether Rab5 may influence CAR protein stability. However, blocking protein synthesis (cycloheximide), lysosome-based degradation (BafA1), or proteasome-based degradation (bortezomib) revealed no differences in CAR surface expression between 19BBz-Rab5 and 19BBz-tEGFR CARTs, regardless of the presence of tumor cells (Figure S3A-S3F). This further indicates that Rab5 maintains CARs on the T cell surface without altering CAR protein levels. Lastly, transcriptional profiling of Rab5 CARTs in the presence and absence of tumor revealed only modest differences (Figure S3G,S3H), suggesting that differences in surface CAR expression are attributable to Rab5 itself, rather than factors regulated by Rab5.

**Figure 3.**
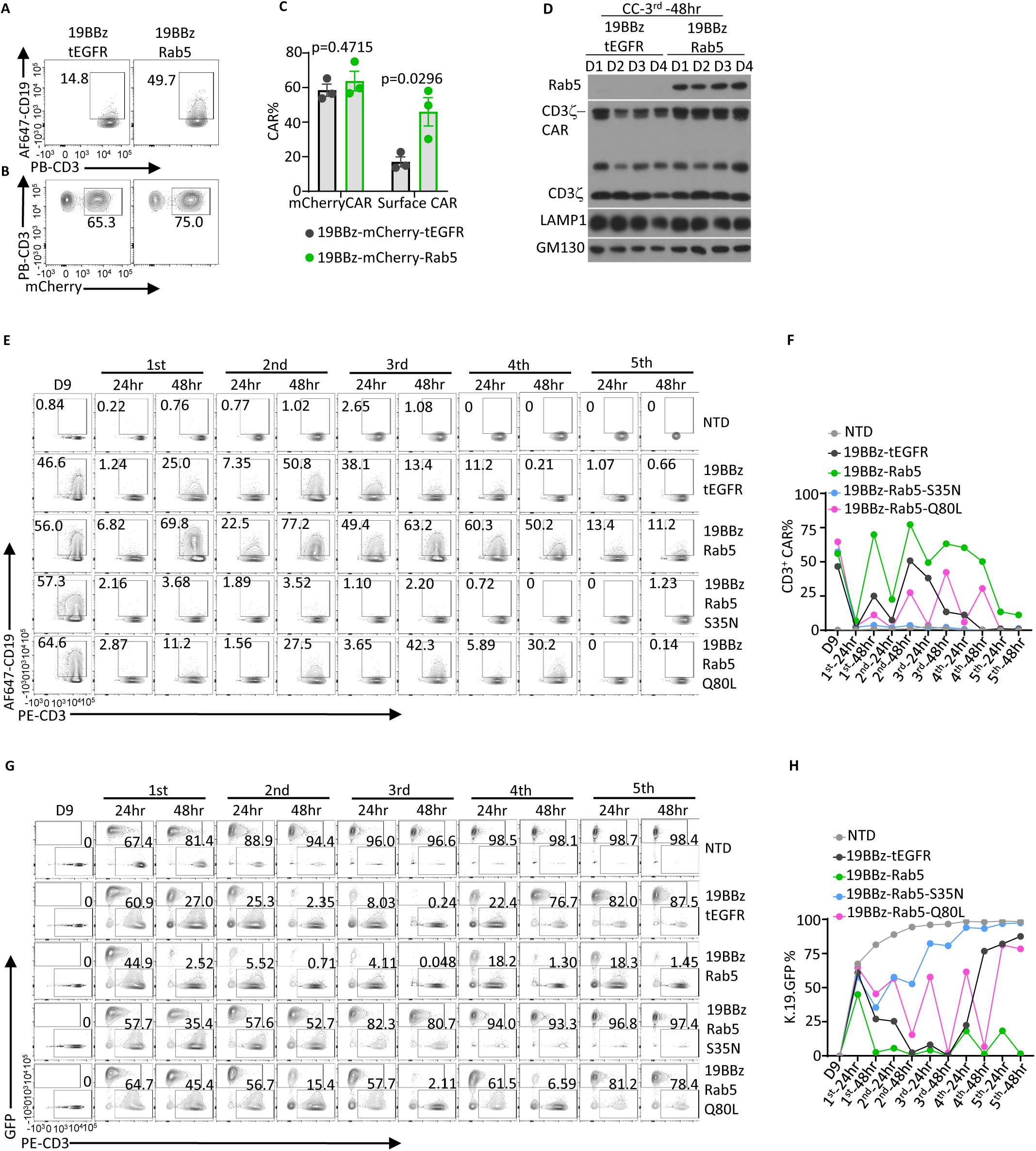
Rab5 maintains surface CAR expression as a GTPase. (A-B). T cells were activated and transduced with 19BBz-mCherry T2A Rab5 or19BBz-mCherry T2A tEGFR and placed into CAE assay. 48 hours after the second time tumor challenge, surface CAR expression was measured by staining with labeled CD19 protein (A) and total CAR expression was determined by measuring mCherry (B). (C) Summary data from three independent experiments measuring surface and total CAR expression. (D). 48 hours after the third tumor challenge in CAE assay, CAR positive T cells were purified by fluorescence-activated cell sorting (FACS). Whole cell lysate from CAR positive cells was used for western blot analysis with indicated protein. Data from four donors (D1-D4) are shown. (E-H). T cells transduced with 19BBz-tEGFR, 19BBz-Rab5, 19BBz-Rab5 S35N, or 19BBz-Rab5 Q80L were placed in the CAE assay. Surface CAR expression was measured by staining with labeled CD19 protein (E, F) and tumor killing was determined by measuring the population of tumor cells expressing GFP (G, H). Experiments shown in (A)-(C), (E)-(H) are representative of three independent experiments. Error bar show mean ± SEM. P value was determined by Students t test. See also Figure S3.

Rab5’s GTPase functions as an on/off switch for its activity^33,34^. We next investigated whether GTPase activity was required for Rab5 to maintain CAR surface expression by introducing a mutation that blocks GTPase function (S35N), and assessing whether a constitutively active form of Rab5 (Q80L)^34^ could maintain even higher levels of CAR expression. Supporting the importance of GTPase function, the S35N variant of Rab5 acted as a dominant negative mutant, causing rapid and permanent loss of surface CAR. This ultimately resulted in CARTs that had significantly impaired tumor-killing function (Figure 3E, 3H). Surprisingly, the constitutively active Q80L Rab5 mutant was less effective at maintaining both surface CAR expression and tumor control than the wildtype Rab5, though it performed better than the dominant negative of Rab5 (Figure 3E, 3H). Together, these findings suggest that a properly regulated Rab5 is essential for maintaining CAR expression and CARTs function.

### Rab5 maintains surface CAR expression by enhancing CAR endocytic recycling

Our results suggest that Rab5 does not enhance CART function through transcriptional or translational mechanisms. Thus, we next investigated how Rab5 overexpression alters the endocytoic rates of CARs expressed on the cell membrane. To measure endocytosis, 19BBz-tEGFR and 19BBz-Rab5 CARTs were first stained with CD19 protein linked to a His tag and AF647 fluorescent dye (AF647-CD19-His) at 4°C. Following incubation at 37°C to induce endocytosis, the cells were stained again using anti-His Tag antibody at 4°C, and the relative fold change of surface CAR molecules was determined (see Figure S4A for methodology). This analysis revealed that 19BBz-Rab5 CARTs exhibited significantly higher endocytic activity compared to 19BBz-tEGFR CARTs (Figure 4A, 4B). During endocytosis, the early endosome is the initial site for surface receptor trafficking^27,35^. Therefore, during the third tumor challenge in CAE assay, we purified early endosomes from 19BBz-mCherry T2A tEGFR and 19BBz-mCherry T2A Rab5 CARTs. In the endosomes from 19BBz-mCherry T2A tEGFR CARTs, we observed natural CD3z and an early endosome marker EEA1. In contrast, in 19BBz-mCherry T2A Rab5 CARTs we observed Rab5, mCherry, CD3z and a larger form of CD3z representing the CAR (Figure 4C). This indicates that Rab5 promotes endocytosis of CAR molecules and retains CARs in the early endosome, protecting them from being captured by tumor cells. To further explore the relationship between surface CAR maintenance and CART-mediated internalization, we pre-treated 19BBz-tEGFR or 19BBz-Rab5 CARTs with Pitstop2 (clathrin-mediated endocytosis inhibitor), Fillipin (clathrin-independent endocytosis inhibitor) or Cytochalasin D (actin polymerization inhibitor) to inhibit CAR internalization, and Plinabulin (tubulin polymerization inhibitor) or Endosidin-2 (endosomal recycling inhibitor) to disrupt CAR recycling to cell membrane surface. After incubation with K.19.GFP cells for additional 24 hours, we observed that inhibitors of clathrin-independent endocytosis (CIE), such as Fillipin and Cytochalasin D, prevented Rab5 from maintaining surface CAR expression in the presence of tumors, whereas the other inhibitors did not affect the ability of Rab5 to maintain surface CAR expression (Figure S4B). This indicates that Rab5 utilizes a CIE pathway to maintain surface CAR expression in CARTs.

**Figure 4.**
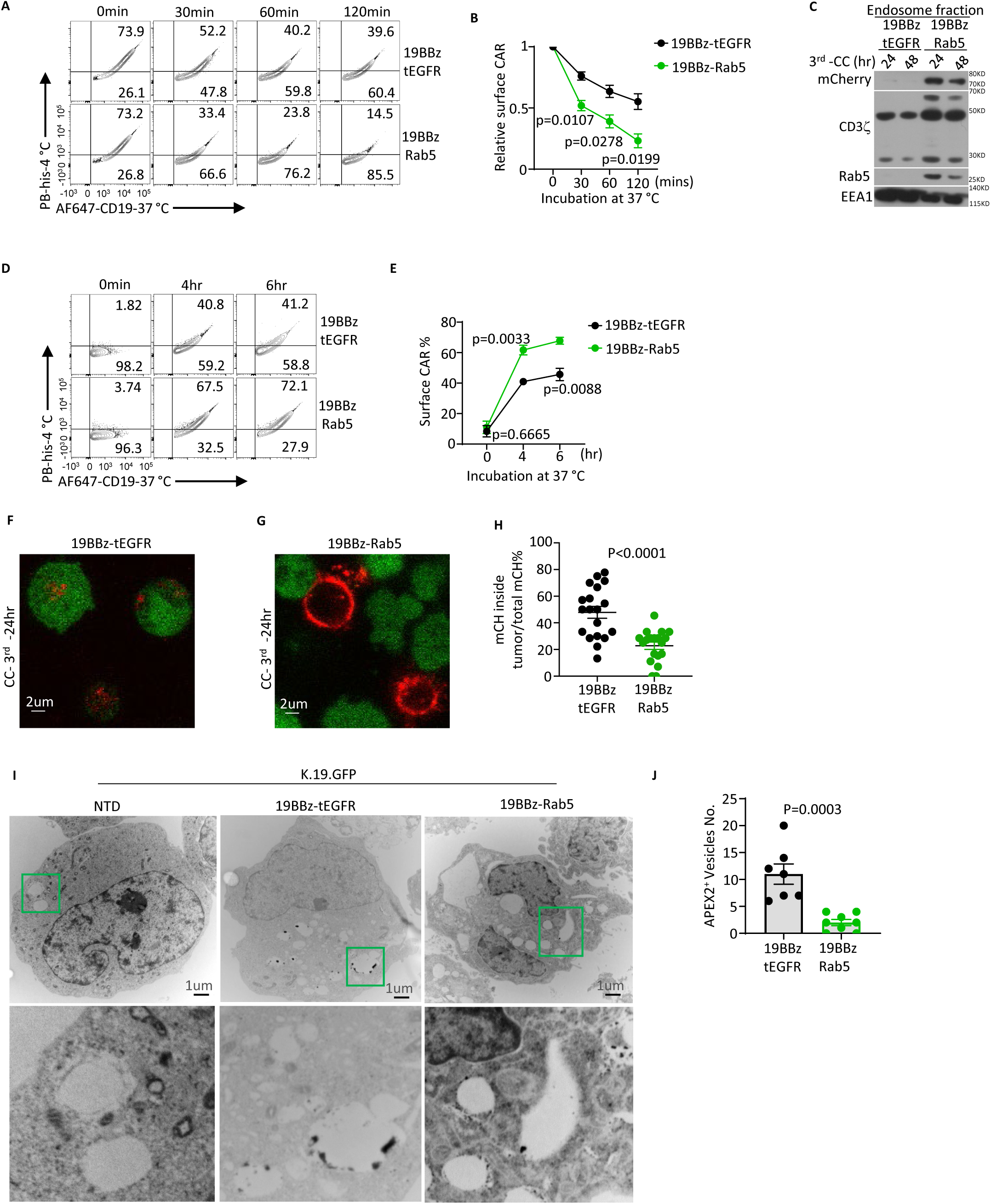
Rab5 increases surface CAR persistence by enhancing endocytic recycling. (A-B). CAR endocytic rate was measured at indicated time point after incubation at 37 °C (A) and presented as relative fold difference between the surface CAR expression of 19BBz-Rab5 and 19BBz-tEGFR CARTs (B). See Figure S4A for detailed workflow. (C). 19BBz-tEGFR and 19BBz-Rab5 CARTs were isolated after 24 hours and 48 hours from the third time tumor challenge of CAE assay. Early endosome fraction was purified and immunoblotting using antibody specific for the indicated proteins was performed. (D-E). CAR recycle rate was measured over 6 hour (D) and percentage of surface CAR difference (E) is performed using 19BBz-tEGFR or 19BBz-Rab5 CARTs. see Figure S4C for detailed workflow. (F-G). Confocal imaging of CAR localization in CARTs and tumor cells. 19BBz-mCherry T2A tEGFR (F) or 19BBz-mCherry T2A Rab5 CARTs (G) (red) co-cultured with K.19.GFP cells (green) for three rounds, 24 hours after that the sub-cellular localization of CAR was measured by confocal microscopy. (H). Quantification of the percentage of mCherry-CAR internalization into GFP-fused tumor cells in (F-G). (I). TME imaging of CAR localization in tumor cells. CAE assay was performed with 19BBz-APEX2-tEGFR or 19BBz-APEX2-Rab5 CARTs with K.19.GFP cells. 24 hours after the third tumor challenge, CARTs were depleted with CD3 beads, the tumor cells were collected for TEM imaging of APEX2-CAR (black) inside of tumor cells. (J) Presentative of the mean number of APEX2-CAR positive vesicles inside of tumor cells in (I). Experiments shown in (A), (C), (D), (F)-(J) are representative of three independent experiments. Error bar show mean ± SEM. P value was determined by Students t test. See also Figure S4.

The early endosome is a sorting endosome where cargos can be directed to the lysosome for degradation or back to the plasma membrane^36^. Our results indicate Rab5-regulated surface CAR expression is not dependent on lysosome mediated degradation (Figure S3B, S3E). To investigate if Rab5 overexpression promotes the endocytic recycling pathway, we performed an assay measuring CAR recycling (Figure S4C) and found 19BBz-Rab5 CARTs displayed much higher recycling rate than 19BBz-tEGFR CARTs (Figure 4D, 4E). This suggests that Rab5 promotes sorting of CARs into endocytic recycling pathway. Next, we assessed whether 19BBz-Rab5 CARTs were protected against “CAR-jacking” by tumor cells. As first shown in Figure1C, we co-cultured mCherry-fused CARTs with K.19.GFP cells and observed high levels of CARs within the tumor cells (Figure 4F). In contrast, 19BBz-Rab5 CARTs retained CAR surface expression, successfully preventing CAR transfer to tumors (Figure 4G, 4H). Likewise, K.19.GFP incubated with 19BBz-Rab5 CARTs captured much fewer APEX2-fused CARs in the intracellular vesicles than those co-cultured with 19BBz-tEGFR CARTs (Figure 4I, 4J). Collectively, these results demonstrated that Rab5 promotes endocytic recycling of CAR, dramatically reducing tumor-mediated CAR loss. This ultimately allows CARTs to retain CARs surface expression and killing capacity during long-term engagement with tumor cells.

### Rab5 maintains CART membrane protrusions

To further explore the effects of persistent Rab5 expression on CARTs, scanning electron microscopy (SEM) was performed to image the surface of CARTs throughout the CAE assay. Interestingly, we observed small membrane protrusions on the surface of CARTs prior to the CAE assay, regardless of Rab5 overexpression. However, 19BBz-Rab5 CARTs exhibited a more of a corrugated membrane compared to 19BBz-tEGFR CARTs (Figure 5A, 5B). After co-culturing with tumor cells for 2 hours, both 19BBz-Rab5 and 19BBz-tEGFR CARTs formed extended protrusions (Figure 5C,5D). Notably, by the third tumor challenge, 19BBz-tEGFR CARTs had lost most of their membrane protrusions, whereas 19BBz-Rab5 CARTs maintained abundant and large protrusions (Figure 5E,5H). A similar phenotype has been described for NK cells where the presence of membrane protrusions correlated with the ability to form immunological synapses and kill targets^37,38^. Our findings suggest that the ability of CARTs to clear tumors directly correlates with their capability to maintain membrane protrusions on the cell surface. Furthermore, Rab5 appears to play a critical role in forming or maintaining these membrane protrusions.

**Figure 5.**
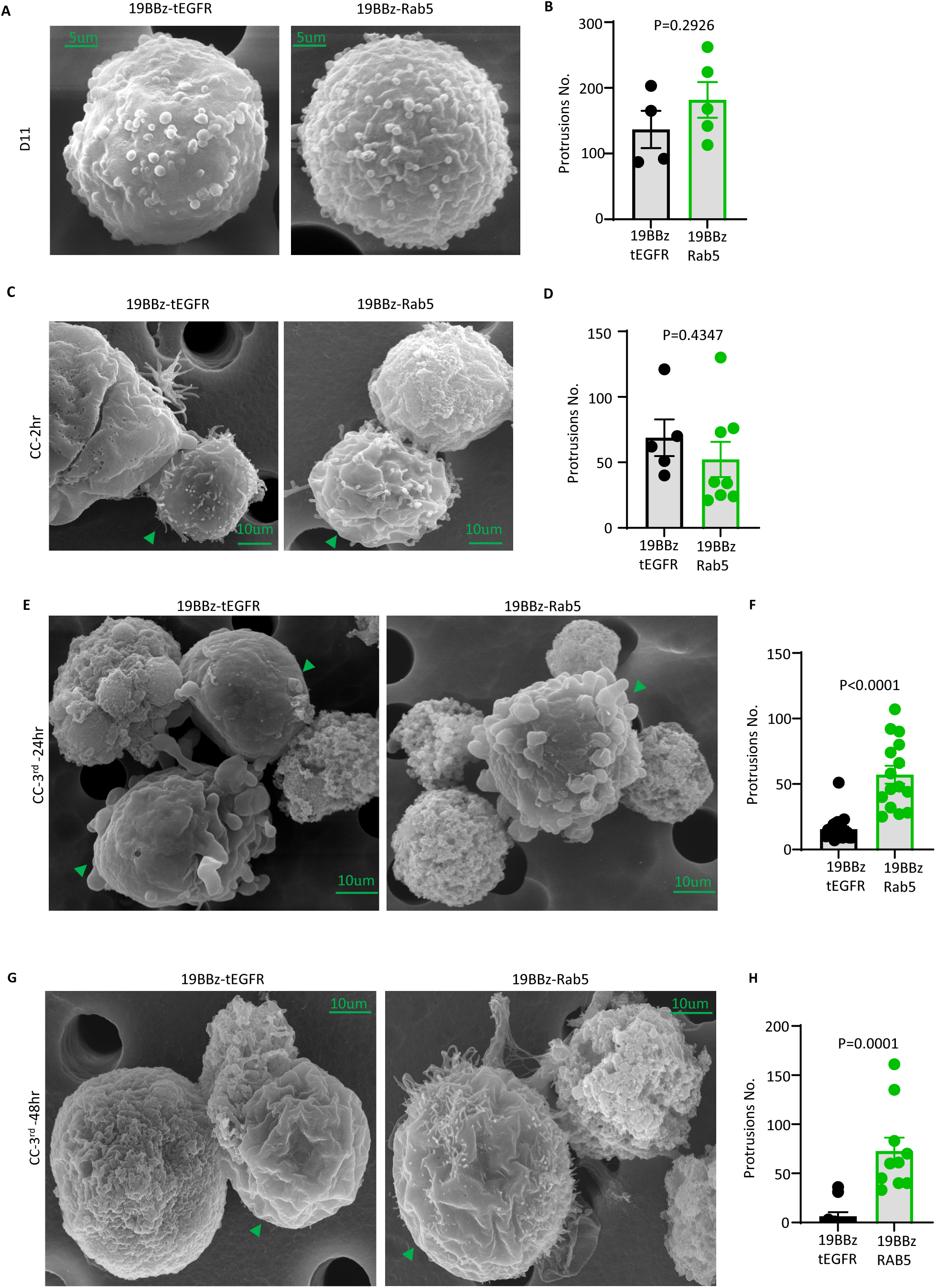
Rab5 maintains CART membrane protrusions. (A-B). Primary T cells were transduced 19BBz-tEGFR or 19BBz-Rab5 vector and cultured for 11 days. Scanning electron microscopy (SEM) was performed with these CARTs. (C-D). SEM imaging of 2-hour co-culture mixture of 19BBz-tEGFR or 19BBz-Rab5 CARTs with K.19.GFP cells. (E-H) CAE assay was performed for three round of tumor re-challenge, samples from 24 hours (E-F) or 48 hours (G, H) after that were imaged by SEM. Summary data was generated by manual counting the number of protrusions on randomly chosen CARTs. One dot represents the number of protrusions on a single CART cell. Data is representative of two independent experiments. Error bar show mean ± SEM. P value was determined by Students t test.

### Rab5 CARTs exhibit superior in vivo antitumor efficacy

We next assessed the *in vivo* efficacy of Rab5-CARTs using well-characterized CD19 CARTs. Utilizing a previously established mouse model, in which K562 cells were injected into the flank of NSG mice to form solid tumors^39^, we observed that 19BBz-Rab5 CARTs cleared tumor rapidly and completely, with all treated mice remaining tumor-free, whereas only two of the five mice treated with 19BBz-tEGFR CARTs were tumor-free after 28 days (Figure 6A-6C and Fig S5A). To further define the interactions between tumors and CARTs *in vivo*, we harvested tumors from mice treated with CARTs for 14 days in a separate experiment. Tumors in mice treated with 19BBz-Rab5 CARTs were significantly smaller (Figure 6D), contained fewer tumor cells (Figure 6E), and had a higher infiltration of CARTs (Figure 6F, 6G) compared to tumors from mice treated with 19BBz-tEGFR CARTs. Also, peripheral blood (PB) isolated from mice treated with19BBz-Rab5 CARTs showed higher CARTs expansion compared to those treated with 19BBz-tEGFR CARTs (Figure 6F, 6H). Importantly, we observed much higher CAR expression in the 19BBz-Rab5 CARTs within the tumors (Figure 6I, 6J), and in the peripheral blood (Figure 6K), confirming that the *in vitro* CAE assay models *in vivo* conditions faithfully. Moreover, tumor cells from mice treated with 19BBz-Rab5 CARTs captured much less CAR molecules compared to those isolated from mice infused with 19BBz-tEGFR CARTs (Figure S5B). Recent clinical data suggest that the presence CD4^+^ CARTs correlate with durable remission^40^. Notably, CD4^+^ 19BBz-Rab5 CARTs appeared to benefit more from Rab5 expression than CD8^+^19BBz-Rab5 CARTs in the tumor (Figure 6L, 6N) and in peripheral blood (Figure 6O, 6P), although all CARTs showed similar CAR expression before injection (Figure S5C). Collectively, these results demonstrate that 19BBz-Rab5 CARTs have superior *in vivo* efficacy, which correlates with the ability of Rab5 to maintain surface CAR expression in the tumor microenvironment.

**Figure 6.**
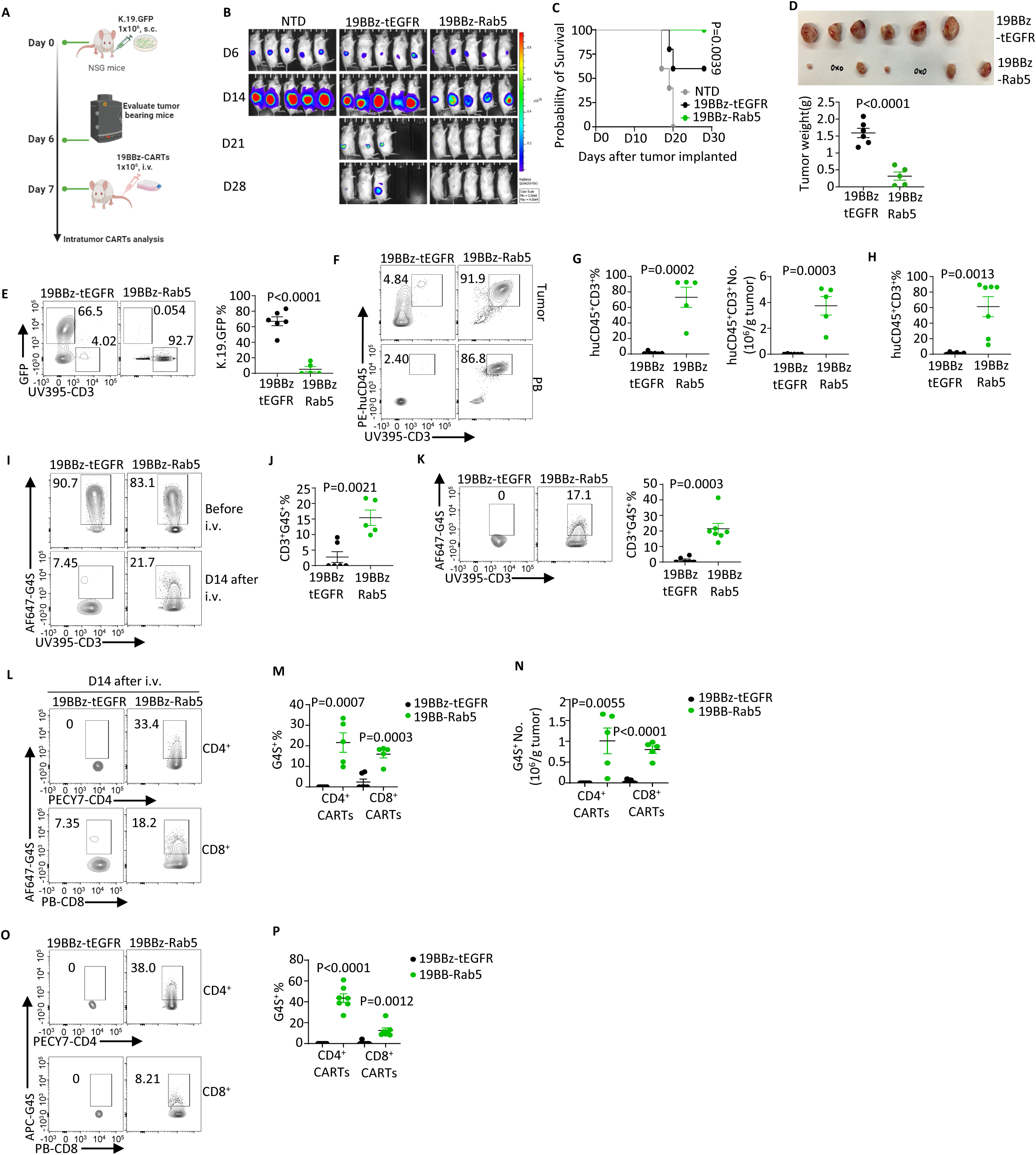
Rab5 CARTs exhibit superior in vivo antitumor efficacy. (A). Experimental design of in vivo tumor challenge mouse model. 1x10^6^ K.19.GFP cells mixed with matrigel were subcutaneously injected (s.c.) to NSG mice on day 0. Tumors were imaged six days later, followed by tail intravenous injection (i.v.) with 1x10^6^ CD19-CAR positive CARTs on day7. (B). Tumor progression was monitored weekly using bioluminescent imaging (BLI) for mice treated with the indicated CARTs. N=5 per group. (C). Survival curve of tumor bearing mice. N=5 per group. (D). In a separate experiment, 14 days after CART infusion, tumor tissues were isolated from mice treated with 19BBz-tEGFR or 19BBz-Rab5 CARTs and weighed. (E). In vivo tumor killing efficacy of 19BBz-tEGFR or 19BBz-Rab5 CARTs was measured by detecting GFP-expressing tumor cells. (F-H). CARTs expansion in tumor tissue (F, G) or in peripheral blood (PB, F, H) from 14 days after indicated CARTs infusion in (D). (I-K). Prior to infusion, 19BBz-tEGFR and 19BBz-Rab5 CARTs were stained for CAR expression using anti-G4S linker antibody (I).14 days after infusion, CARTs from tumors (I, J) and peripheral blood (K) were tested for surface CAR expression. (L-N). Comparison of surface CAR expression level (L, M) and absolute CARTs number (N) of CD4^+^ and CD8^+^ CARTs isolated from tumors. (O-P). Comparison of surface CAR expression on CD4^+^ and CD8^+^ CARTs from PB after 14 days CARTs infusion. Data shown in (B, C), is representative of three independent experiments using T cells from different donors, N=5. (D)-(P), the data is representative of two independent experiments. For tumor tissues, N=6 for 19BBz-tEGFR CARTs, N=5 for 19BBz-Rab5 CARTs; For PB samples, N=6 for 19BBz-tEGFR CARTs, N=7 for 19BBz-Rab5 CARTs. In (C), survival curves were compared using the log-rank Mantel–Cox test. Error bar show mean± SEM. P value was determined by Students t test. See also Figure S5.

### Rab5 expression enhances the ability of CARTs to control mesothelin-expressing tumors

While the above results mimic a solid cancer model using CD19-specific CARTs, we next investigated whether Rab5 could enhance the efficacy of CART therapy for a true solid tumor, mesothelioma. The M5 CAR (MESO-CAR) targets mesothelin and has shown promising activity in a Phase I clinical trial^41^ but durable cures have not occurred yet. We infused EMMESO tumors into NSG mice and treated these mice with non-transduced T cells (NTD), MESOBBz-tEGFR or MESOBBz-Rab5 CARTs (Figure 7A). Compared to MESOBBz-tEGFR CARTs, MESOBBz-Rab5 CARTs cleared tumors much more rapidly in all tumor-bearing mice but one, without causing significant body weight loss (Figure 7B and Figure S6A). In separate experiments, we isolated tumors and spleens 21 days tumors post-CART infusion. Tumors from mice treated with MESOBBz-Rab5 CARTs were significantly smaller than those treated with MESOBBz-tEGFR CARTs (Figure 7C, 7D). Additionally, more tumor-infiltrating CARTs were observed in MESOBBz-Rab5 CART-treated tumors (Figure 7E,7F). Rab5 expression enhanced surface CAR expression both on intratumor CARTs (Figure 7G,7H) and splenic CARTs (Figure 7I), and it also reduced the uptake of CARs into tumor cells (Figure S6B). Moreover, CD4 T cells were more significantly impaired by “CAR-jacking” than CD8 T cells. However, Rab5 restored CAR surface expression on CD4^+^ CARTs both in the tumor microenvironment (Figure 7J,7K) and in the spleen (Figure 7L). A similar phenotype was observed from intratumor CD8^+^ T cells (Figure S6C, S6D) and splenic CD8^+^ T cells (Figure S6E). Collectively, these data confirm that engineering CART with Rab5 significantly promotes their therapeutic efficacy against solid tumors *in vivo*.

**Figure 7.**
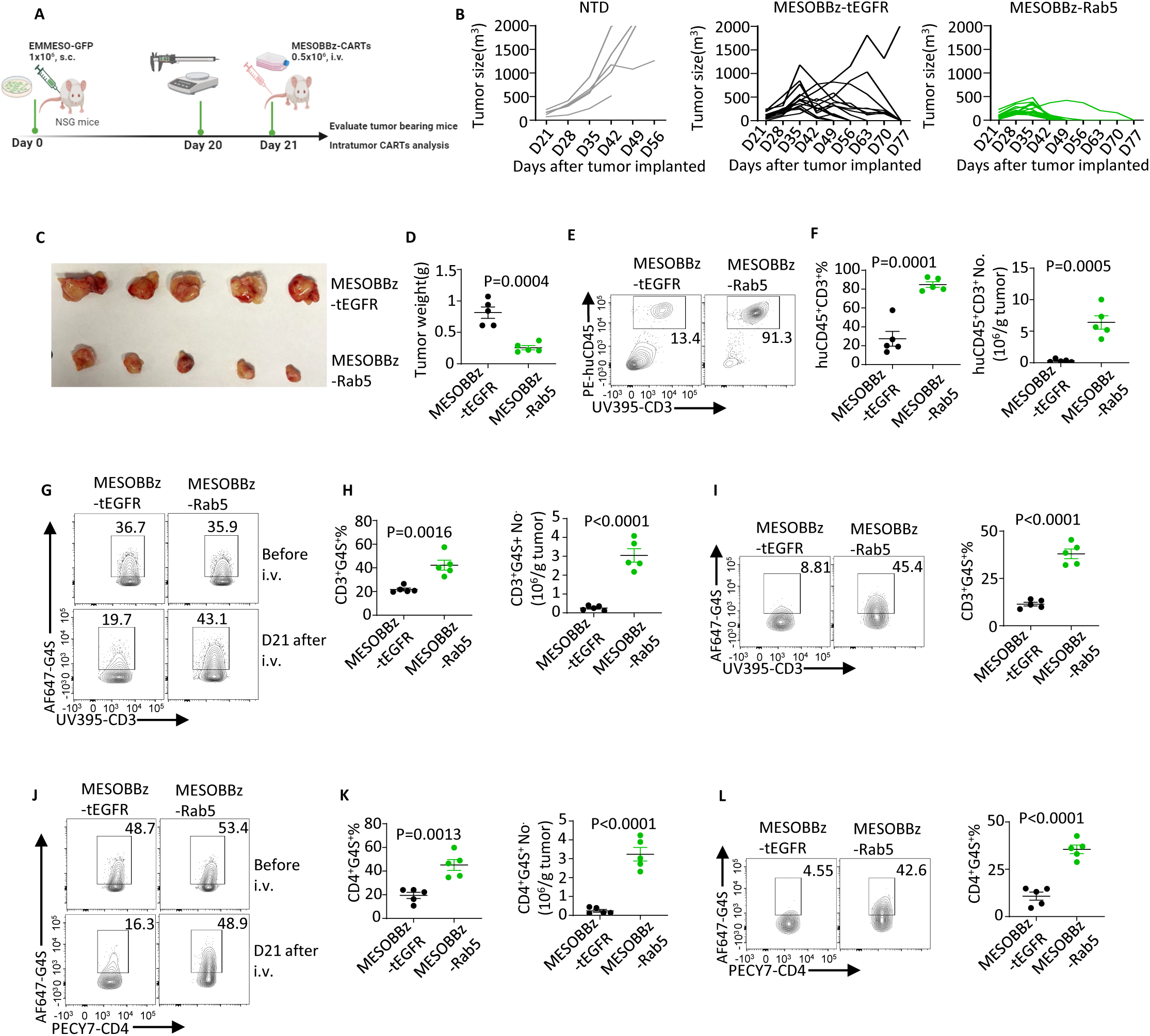
Rab5 expression augments the ability of CAR T cells to control solid tumors. (A). Schematic of EMEMSO tumor mouse model. 1x10^6^ EMMESO-GFP cells were subcutaneously injected (s.c.) to NSG mice. Tumor size reached around 150mm^3^ after 20 days, 0.5x10^6^ non-transduced (NTD), MESOBBz-tEGFR or MESOBBz-Rab5 CARTs were infused i.v. on day 21. (B). Individual tumor burden was measured by calipers weekly. N=5 for mice treated with NTD T cells, N=15 for mice treated with MESOBBz-tEGFR or MESOBBz-Rab5 CARTs. (C-D). Imaged (C) and weight (D) of tumor tissues isolated on day42 from MESOBBz-tEGFR or MESOBBz-Rab5 CARTs treated mice separately. (E-F). Tumor infiltrating CARTs from (C) were stained with anti-huCD45 and anti-huCD3. (G-I). Surface MESOBBz-tEGFR and MESOBBz-Rab5 CAR expression before (G) or 21 days after CARTs infusion were detected by anti-G4S linker antibody staining with tumor samples (G, H) or spleen samples (I). (J-K). Surface MESO-CAR expression on CD4^+^ CARTs before (J) or after infusion were isolated from tumor (J, K) or spleen (L). Data shown in (B) is representative of three independent experiments. Data shown in (C)-(L) is representative of two independent experiments with different donors, N=5 per group. Error bar show mean± SEM. P value was determined by Students t test. See also Figure S6.

## DISCUSSION

CARs are synthetic constructs composed of domains from various proteins, designed to effectively redirect T cells^42^. These domains typically confer supraphysiologic properties to CARTs. For instance, incorporating costimulatory domains such as 4-1BB and CD28 to the CD3 zeta chain enables CARTs to receive both signal one and signal two through a single receptor^43,44^. However, some of these properties may interfere with, rather than promote, CART activity. Our studies uncover one such instance. The TCR/CD3 complex is a remarkable immunosensor capable of distinguishing between self and non-self through low affinity interactions, subsequently initiating a signaling cascade that promotes T cell expansion and effector functions^45^. Critical to maintaining an orchestrated signaling cascade is prevention of new initiation events and overstimulation, which can drive inflammatory damage and autoimmunity^46^. As such, multiple mechanisms ensure TCR downregulation after T cell activation, including ubiquitination-mediated degradation of the CD3 zeta chain^47^, and Cbl family-mediated clearance of surface-expressed TCR^48^. Furthermore, a more recently described mechanism, TCR ectocyotosis^16^, has significant implications for CART biology. In this process, TCRs that have productively engaged with pMHC are targeted. CD3 signaling temporally inactivates local endocytosis, which can allow the pMHC-presenting cell to capture activated TCRs. This effectively limits additional TCR activation by removing these TCRs from the T cell surface. Since only a minor fraction of TCRs are lost this way due to the scarcity of cognate pMHC on the target cell surface, the T cell can maintain TCR expression without issue. However, CARTs differ markedly from non-engineered T cells in this context. Up to 100,000 of CAR molecules on the T cell surface^49^ can interact with up to 100,000 molecules on the tumor surface^50^. Since all these interacting CARs have the potential to become activated, the above ectocytosis process is capable of permanently removing a large fraction of CARs from the T cell surface. Our studies show that impressive numbers of CARs are captured by tumor cells. Thus, while T cells have some capacity to maintain CAR expression, repeated tumor engagements deplete this reserve and leave the CART with insufficient CAR levels to kill tumor, resulting in significant CART dysfunction. While it is difficult to determine how much CAR expression is lost due to “CAR-jacking” (versus other mechanisms of CAR loss), the ability of Rab5-mediated endocytosis to restore high levels of CAR expression suggest that ‘‘CAR-jacking’’ is a major reason for the loss of CAR on the CART surface.

Endocytosis encompasses at least five distinct pathways and involves over 60 proteins that work collaboratively to regulate the lipid and protein composition of the plasma membrane^51^. Our studies indicate that Rab5 expression is necessary to maintain CIE, and that is the primary mechanism through which Rab5 maintains surface CAR expression. While immune cells such as dendritic cells and macrophages exhibit significantly higher endocytic activity than T cells, endocytosis does play a crucial role in T cell biology^52^. Despite this, we observed only a transient elevation of Rab5 after T cell activation. This low level of Rab5 in T cells is likely required to allow for the precise regulation of TCR surface expression following activation, as described above. The importance of Rab5 to CART biology is underscored by our finding that introducing a GTPase-mutated, dominant-negative version of Rab5 led to rapid loss of CAR expression on the cell surface. Our studies also highlight that overexpression of Rab5 appears to sufficiently augment endocytic activity, suggesting that Rab5 functions as the rate-limiting component of endocytosis in T cells. Notably, expression profiling of CARTs with and without Rab5 revealed no global changes in gene expression, and, with the exception of increased Rab5 expression, only modest alterations in the expression of a small set of genes. This suggests that Rab5 primarily enhances the endocytic process without affecting established signal transduction pathways.

Previous studies have shown overexpression of various Rab GTPases often induces specialized cellular phenotypes, such as changes in cell morphology, cytoskeletal rearrangements, and even organelle structure^22,53^. Our studies suggest that Rab5 plays a critical role in maintaining cell membrane protrusions. In T cells, these finger-like protrusions, also known as ‘microvilli’, play a crucial role in scanning antigens through TCRs, allowing TCRs to respond to pMHC in a sensitive and rapid manner^54^, forming T cell-target contacts^55^, facilitating the endocytosis of the IL-2 receptor^56^, and activating T cells^57^. The shedding of TCRs in microvilli represents another mechanism for TCR downregulation following T cell activation^58^. The role of microvilli in CARTs is less clear, given the higher affinity of CAR/target interactions and the abundance of CARs on the surface. Nonetheless, we observed a strong correlation between the loss of membrane protrusions and downregulated CAR surface expression. Our studies revealed an unexpected relationship between microvilli, endocytosis, CAR expression, and cytolytic potential. It is intriguing to consider the sequence of these events, their interdependence, and how this relationship can be further harnessed to optimize and innovate CART therapy.

In summary, the vast number of tumor cells that must be eliminated to achieve a durable cancer cure presents a formidable challenge for current CART strategies. Here, we have identified an additional obstacle, termed ‘‘CAR-jacking’’, where repeated tumor encounters result in capturing of CAR molecules by the tumor, thereby impairing the ability of CARTs to maintain CAR expression on their surface. Overexpression of Rab5 overcomes this defect and greatly potentiates CART activity, suggesting that Rab5 overexpression could benefit all CART therapies.

## LIMITATIONS OF THE STUDY

While immunocompromised NSG models are commonly used to assess CART therapy efficacy, models with human tumors interacting with the human immune system are necessary. Hence, the antitumor efficacy of Rab5-CARTs should be evaluated in future clinical trials. One potential side-effect of Rab5 overexpression might be the hyperactivity of the TCR within the CARTs, which could be problematic once tumor is cleared. We would like to note that inserting CAR and Rab5 into the TCR locus^59^ could eliminate this risk and provide an additional rationale for disrupting TCR expression in CARTs.

## ACKNOWLEDGMENTS

Research supported by U19AI149680, UM1AI164570, and sponsor research from Tmunity/Kite Pharma. We thank Max Eldabbas, Emileigh Maddox, Lynn Chen and Miranda Wang of the Human Immunology Core (RRID: SCR_022380) at the Perelman School of Medicine at the University of Pennsylvania for providing purified human T cells; Biao Zuo and Inna Martynyuk of Electron Microscopy Resource Lab at the Perelman School of Medicine at the University of Pennsylvania(SCR_022375) for providing advice and assistance of TEM preparation and imaging ; Yuri Veklich of CDB Microscopy Core (RRID: SCR_022373) at the Perelman School of Medicine at the University of Pennsylvania for processing and imaging of SEM samples; Stem Cell and Xenograft Core (RRID: SCR_010035) at the Perelman School of Medicine at the University of Pennsylvania for providing and housing NSG mice; Flow Cytometry and Cell Sorting Facility (RRID: SCR_ 022376) at the Perelman School of Medicine at the University of Pennsylvania for cell sorting and maintaining flow cytometry daily. We also thank Drs. Carl H. June (M5-BBz CAR vector), Steve Albelda (EMMESO), David Barrett, Gavin Ellis, Divanshu Shukla, Sriram Srivatsa, Yuqi Zhou and John Wherry for reagents and helpful advice.

## AUTHOR CONTRIBUTIONS

MG and JLR designed the studies, analyzed, and interpreted the data, and wrote the manuscript. MG, EJC. KAR, and DPN performed experiments and analyzed data. KAR edited the manuscript. All authors approved the final version of the manuscript.

## RESOURCE AVAILABILITY

### Lead contact

Further information and requests for resources and reagents should be directed to and will be fulfilled by the Lead Contact, Dr. James L. Riley.

### Materials availability

All resources and materials reported in this paper will be shared by the lead contact upon request.

### Data and Code availability

- Bulk-RNA seq data has been deposited at GEO: GSE269885 is publicly available as of the date of publication.
- Any additional information required to reanalyze the data reported in this paper is available from the lead contact upon request.

## EXPERIMENTAL MODEL AND SUBJECT DETAILS

### Cell lines

Human myelogenous leukemia cell line K562 was transduced with lentiviral vector of pTRPE-huCD19-CBG-GFP minigenes (K.19.GFP), single-cell clones were sorted, and a clone that exhibited stable expression was selected as previously described^60^. Human EM-Meso-GFP-Luc cell line (EMMESO-GFP)^61^ was a kind gift from Dr. Steven M. Albelda laboratory (University of Pennsylvania). K.19.GFP and EMMESO-GFP cells were cultured in R10 media consisting of RPMI-1640 (Gibco) with 10% fetal bovine serum (Avantor Seradigm), 1% HEPES (Gibco), 1% GlutaMAX^TM^ (Gibco), and 1% penicillin/streptomycin (Gibco). All cell lines were regularly tested for free of mycoplasma contamination.

### Mice studies

Animal experiments were performed according to protocols approved by the Institutional Animal Care and Use Committee (IACUC) of the University of Pennsylvania. Six-to twelve-week-old female and male nonobese diabetic (NOD)/severe combined immunodeficient (SCID)/γ-chain^−/−^ (NSG) mice were obtained from the Stem Cell and Xenograft Core of the Abramson Cancer Center at the University of Pennsylvania. Mice were maintained in pathogen-free conditions. Mice health status was routinely checked by qualified veterinarians.

### Human samples

De-identified healthy donor primary T lymphocytes were provided by the Human Immunology Core at the University of Pennsylvania.

## METHOD DETAILS

### Viral vector construction

All lentiviral constructs were cloned into the transfer vector pTRPE. The anti-huCD19 Rab5 CAR plasmid was constructed by obtaining a gblock (IDT) containing the coding sequence of Rab5 (GenBank U18420.1) with AvrII and SalI restriction sites. The gblock was inserted it into the pTRPE backbone, containing the huCD19,4-1BB or CD28 and CD3 zeta signaling domain^62^. The anti-MESO (M5) CAR lentiviral construct^28^ was a gift from Dr. Carl H. June laboratory (University of Pennsylvania). The M5 scFv sequence was digested with NheI and XmaI restriction enzymes and inserted into 19BBz-EGFRt and 19BBz-Rab5 pTRPE vectors, replacing the anti-huCD19 scFv sequence. All final constructs were verified using restriction digests and whole plasmid sequencing via Plasmidsaurus.

### Lentiviral vector production

HEK293T cells were transfected with lentiviral CAR transfer plasmid and packaging plasmids using Lipofectamine 2000 (Invitrogen) following the manufacturer’s protocol. Lentiviral supernatants were collected 24- and 48-hour post-transfection and concentrated using high-speed ultracentrifugation as previously described^44^. The resulting concentrated lentivirus batches were resuspended in cold R10 media and stored at -80℃.

### CAR T cells generation

CARTs were generated as previously described ^28^. Briefly, primary human T cells were maintained in Cell Therapy Systems (CTS), OpTmizer T Cell Expansion Serum Free Medium (Thermo Fisher Scientific) supplemented with CTS OpTmizer T cells Expansion Supplement (Thermo Fisher Scientific), 1% GlutaMAX^TM^ (Gibco), 1% penicillin/streptomycin (Gibco), and 1%HEPES (Gibco). CD4^+^ and CD8^+^ human T cells mixed at 1: 1 ratio and plated at 1x10^6^ total T cells/ml were activated with huCD3/CD28 Dynabeads (Gibco) at a 3: 1 bead-to-cell ratio.

After approximately 24-hours, T cells were transduced with lentiviral vectors by adding the virus supernatant directly to T cell cultures. At day 3, beads were removed from the cell culture. For the remainder of the culture, cells were counted every other day and maintained at 0.5x10^6^ cells/ml.

### CAR T cell in vitro rechallenge model

A total of 2x10^6^ K.19.GFP cells or EMMESO cells were co-cultured with 1x10^6^ CAR-positive T cells in 1 ml OpTmizer T Cell Expansion Medium in a 12-well plate. Every 24 hours, half of the supernatant was isolated for flow cytometry analysis of surface CAR expression and tumor fluorescence. Fresh culture medium (0.5 ml) was then added to the remaining cells. Every 48 hours, surface CAR expression and tumor cell survival were monitored as above. Subsequently, an additional 2x10^6^ tumor cells resuspended in 0.5 ml fresh medium were added to the ongoing co-cultures. This process was repeated up to 10 times.

### Flow cytometry

CD19 CAR expression was assessed using AF647-labeled CD19 protein (0.5 µg/ml) or AF647-G4S linker antibody (1:400). MESO CAR expression was assessed using AF647-labeled MSLN protein (1 µg/ml) for in vitro assays, or AF647-G4S linker antibody (1:400) for in vivo detection. For staining of other surface markers, cells were incubated with a 1:200 dilution of antibodies at 4°C for 30 minutes in FACS buffer (PBS + 0.5% FBS). For intracellular staining, cells were first fixed and permeabilized with BD Cytofix/Cytoperm ^TM^ solution (BD Biosciences) at 4°C for 20 minutes, followed by staining at 4°C for 30 minutes in the dark. Flow cytometry data were acquired using BD LSRFortessa (BD Biosciences) and analyzed with FlowJo software (BD Biosciences). The flow cytometry antibody information is summarized in the Key Resources Table.

### Western blotting and Early endosome fraction isolation

T cells were lysed with 1x Lysis Buffer (CST) containing Protease Inhibitor Cocktail (Thermo Scientific) for 10 minutes on ice. Lysed cells were then centrifuged at 12,000 rpm for 10 minutes at 4°C and the supernatant was collected for immunoblot assays. For analysis of CARs in endosomes, early endosomes from T cells were isolated using the Trident Endosome Isolation Kit (GeneTex) according to the manufacturer’s instructions. Briefly, T cells were first purified from the tumor rechallenge co-cultures using the Dynabeads™ Untouched™ Human T Cells Kit (Invitrogen). After washing with cold PBS, the cell pellet was prepared for endosome isolation following the manufacturer’s protocol. All procedures were performed on ice or at 4°C. The early endosome pellets were resuspended in 80 µl of 1x lysis buffer (CST) for immunoblotting analysis. Antibodies against the indicated proteins are summarized in the Key Resources Table.

### Confocal Imaging

For detecting CARs internalized into tumor cells, mCherry-fused CARTs were incubated with GFP-fused K562 cells at 1:1 ratio of total 1x10^6^ cells for the indicated amount of time. Cells were plated on the glass bottom dishes (ibidi) prior to imaging and live imaged at 37°C using a heated stage on the Zeiss 710 inverted confocal microscope. Images were acquired at 488- and 594-nm excitation with 63x oil immersion objective. Captured images were analyzed with ImageJ software (NIH).

### Transmission Electron Microscopy (TEM)

For transmission electron microscopy (TEM) 1x10^6^ APEX2-fused CARTs were co-cultured with 2x10^6^ K.19.GFP cells for the indicated period. CART and tumor populations were analyzed individually. To facilitate this, CARTs were separated using the Dynabeads™ CD3 kit (Invitrogen) according to the manufacturer’s instructions. The CD3 negative (tumor cells) population was retained for analysis. Purified cells were fixed with 2% paraformaldehyde solution and 2.5% glutaraldehyde solution at 4°C for at least 2 hours. Samples were then prepared for Diaminobenzidine (DAB) staining as previously described ^16^. Briefly, fixed cells were washed with pre-cold 0.1M sodium cacodylate rinsed in 0.5M Tris-HCl buffer (pH 7.6) and incubated in DAB reaction solution (0.75 mg/ml DAB dissolved in HCl and 0.3% v/v hydrogen peroxide in Tris-HCl buffer) for 20 minutes at 4°C in the dark. Samples were washed in Tris-HCl buffer followed by 0.1M sodium cacodylate, and then post-fixed in 0.1% osmium tetroxide for 1 hour at 4°C. After washing with pre-cold dH2O, samples were stained with 0.5% aqueous uranyl acetate overnight at 4°C. The next day, samples were dehydrated with graded concentrations of ethanol (50%, 70%, 80%, 90%, 100%, 100%) and embedded in 1:1 and subsequently pure EPON resin overnight at room temperature. Ultrathin sections were prepared by the Electron Microscopy Resource Lab (University of Pennsylvania). Sections were imaged with a JEOL JEM 1010 transmission electron microscope.

### Scanning Electron Microscopy (SEM)

Samples were prepared from 0.2 x 10^6^ CARTs at the resting state or from a cell mixture including 0.2 x 10^6^ CARTs and 0.4 x 10^6^ K.19.GFP cells, incubated for the indicated times. Scanning electron microscope experiments were carried out at CDB Microscopy Core (University of Pennsylvania). Briefly, samples were washed three times with 50mM Na-cacodylate buffer and fixed overnight with 2.5% glutaraldehyde in 50mM Na-cacodylate buffer (pH 7.3) at 4°C. Then samples were dehydrated with a graded series of ethanol concentrations with (70%, 80%, 90%, 100%), for 10mins each. Dehydration in 100% ethanol was performed three times. After dehydration, samples were incubated for 20min in 50% HMDS in ethanol followed by three changes of 100% HMDS (Sigma-Aldrich). After HMDS treatment, samples were removed for overnight air-drying as described previously ^63^. Samples were mounted on stubs and sputter coated with gold palladium. Specimens were observed and photographed using a Quanta 250 FEG scanning electron microscope (FEI, Hillsboro, OR, USA) at 10 kV accelerating voltage. The mean number of membrane protrusions was obtained by analyzing the number of protrusions in a single cell.

### Bulk RNA-seq and Data analysis

Human T cells from four different healthy donors were activated with CD3/28 coated beads and transduced with either 19BBz-tEGFR or 19BBz-Rab5 vectors. To profile baseline differences between these two populations, CARTs expanded *in vitro* for 9 days were collected for analysis. To profile differences under repeated tumor challenges, beginning at day 9, CARTs were co-cultured with K.19.GFP cells at a 1:2 ratio (CARTs:Targets). Every 2 days, target cells were replenished to rechallenge CARTs. 48-hours after the third round of target re-challenge, CD3^+^CAR^+^ cells were enriched. Dead cells were removed using the Dead Cell Removal Kit (Miltenyi Biotec). T cells were isolated from target K562 cells using the Dynabeads™ Untouched™ Human T Cells Kit (Invitrogen). Surface CAR-positive cells were sorted by flow cytometry. A portion of each sample was reserved to test Rab5 overexpression via immunoblotting. For RNA-sequencing analysis, another portion (1x10^6^ cells per sample) was flash-frozen in liquid nitrogen and sent to BGI Genomics for sample QC, mRNA isolation, RNA library preparation and sequencing. To address potential variability in genes with low read count, filtering was performed to omit genes in which more than 4 samples had a read count <10. Genes with an adjusted p-value <0.05 were considered significant, and those with fold changes of >3 were defined as differentially expressed genes (DEGs) for the present study.

### Xenograft Mouse model

NSG mice were purchased from and housed in the Stem Cell and Xenograft Core of the Abramson Cancer Center (University of Pennsylvania), under pathogen-free conditions. Animal studies were approved by the University of Pennsylvania Institutional Animal Care and Use Committee. For the CD19-CART-K562 mouse model, NSG mice were subcutaneously (s.c.) injected with 1 x 10^6^ K.19.GFP cells suspended in 100 µl Matrigel (CORNING): PBS (1:1) into the right flank. Six days after tumor implantation, 1 x 10^6^ CAR-positive CD19-CARTs from 10 days of in vitro culture were intravenously (i.v.) injected into tumor-bearing mice. Tumor progression was monitored by bioluminescence imaging every 7 days. Body weight and survival conditions were checked weekly. For the EMMESO mouse model, NSG mice were s.c. challenged with 1 x 10^6^ EMMESO cells in the right flank. When the mean tumor volume reached 150 mm^3^, mice were i.v. treated with 0.5 x 10^6^ MESO-CAR-positive CARTs. Tumor volumes were calculated using caliper measurements. Tumor growth and body weight were assessed weekly. At the indicated days after CARTs infusion, mice were sacrificed, and tumors were harvested and digested with Tumor Dissociation Kit (Miltenyi Biotec) following the manufacturer’s protocol. Tumor infiltrating lymphocytes were processed with Lymphocyte Cell Separation Medium (Cedarlane) by centrifugation (1200g, 20mins). Spleen samples from tumor-bearing mice were prepared by mashing them through 100-µm filters (Falcon), followed by red blood cell lysis with ACK Lysis Buffer (Lonza). Peripheral blood samples were also treated with ACK Lysis Buffer for 10 minutes on ice. T cells from the above suspensions were identified by flow cytometry with anti-mouseCD45^-^, anti-humanCD45^+^ and anti-humanCD3^+^ antibodies.

## QUANTIFICATION AND STATISTICAL ANALYSIS

Significance of in vivo tumor bearing mice survival data was calculated using the log-rank Mantel-Cox test. Statistical analyses on flow cytometry, confocal imaging, TEM and SEM quantification were determined by unpaired two tailed Student’s t test. All values and error bars are presented as mean ± SEM. All statistical tests were performed on GraphPad 9.

