## Supplemental Figure and Figure legend for "Rab5 Overcomes CAR T Cell Dysfunction Induced by Tumor-Mediated CAR Capture"

### SFig1. CAR expression is lost after repetitive tumor challenge

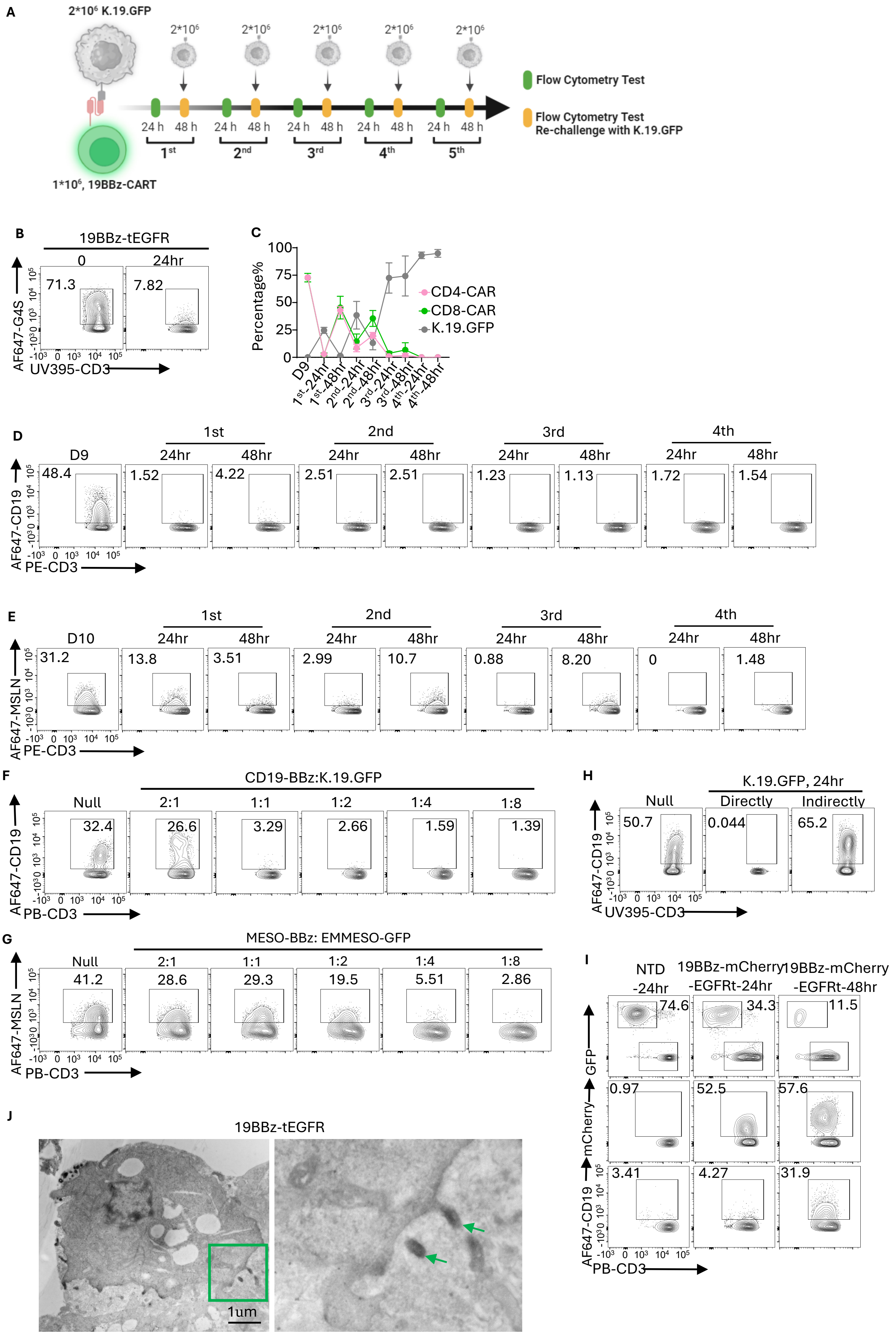

### SFig2. Rab5 enhances surface CAR persistence and CART function

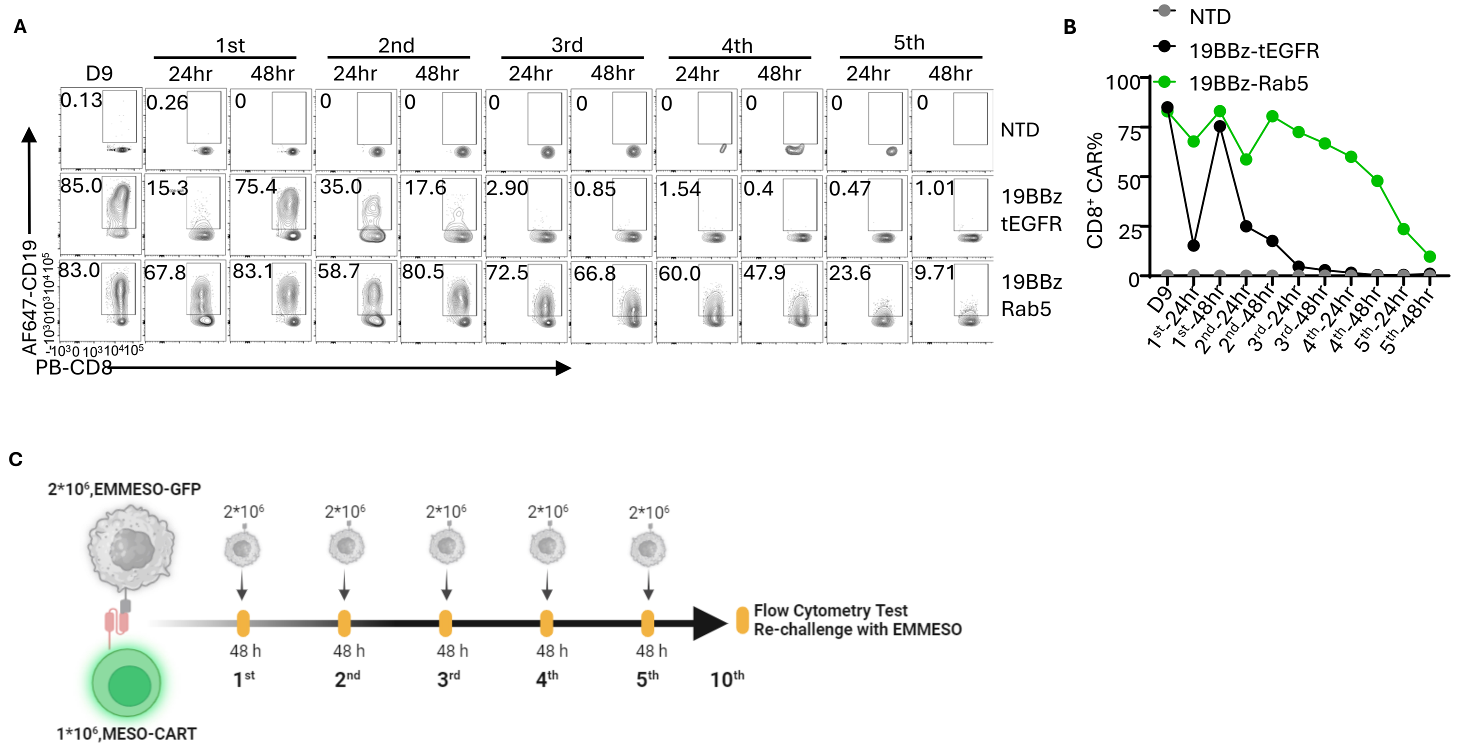

### SFig3. Rab5 has no effect on CAR molecular stability neither transcriptional level

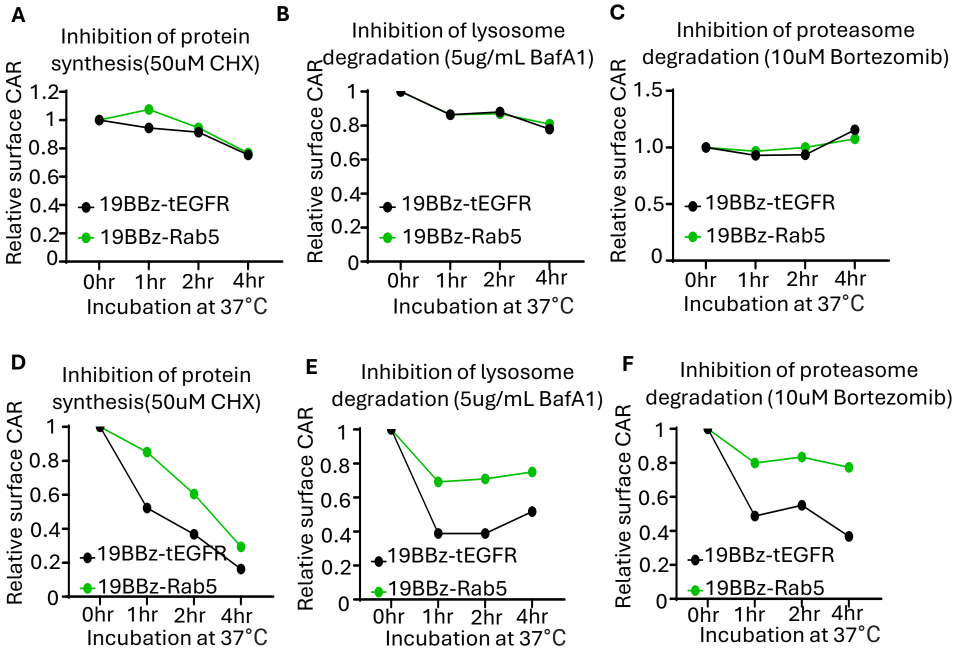


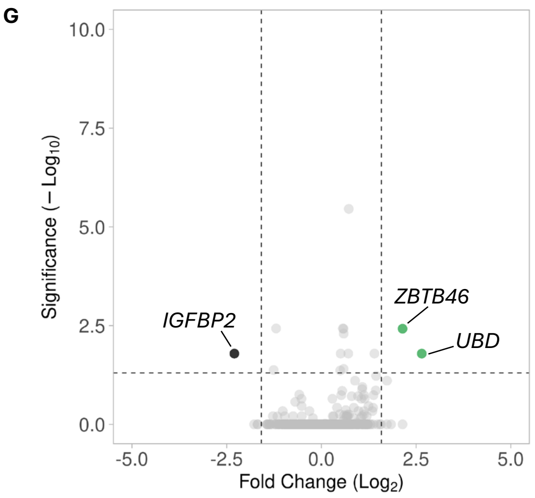


| Upregulated Genes | | |
|---|---|---|
| Gene | Log2FC | adjusted p-value |
| *UBD* | 2.649465831 | 0.039554962 |
| *ZBTB46* | 2.141944042 | 0.00379403 |

| Downregulated Genes | | |
|---|---|---|
| Gene | Log2FC | adjusted p-value |
| *IGFBP2* | -2.2972581 | 0.01619918 |

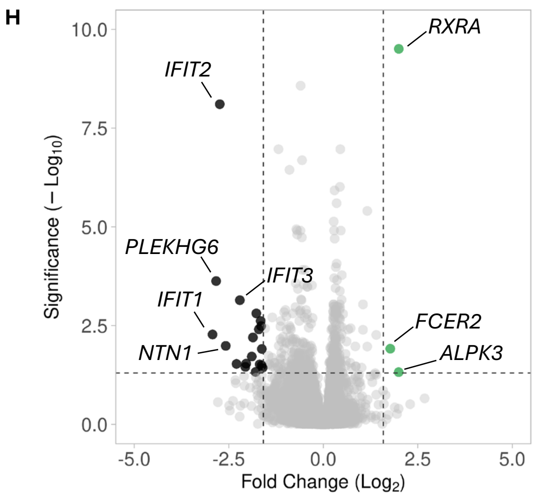


| Upregulated Genes | | |
|---|---|---|
| Gene | Log2FC | adjusted p-value |
| *RXRA* | 1.99609905 | 2.66354E-09 |
| *ALPK3* | 1.99415508 | 0.048314316 |
| *FCER2* | 1.77138009 | 0.012182446 |

| Downregulated Genes | | |
|---|---|---|
| Gene | Log2FC | adjusted p-value |
| *CLIC6* | -1.6065865 | 0.037098 |
| *CRTAM* | -1.6252835 | 0.013004879 |
| *TSGA10* | -1.6311532 | 0.032327313 |
| *BVES* | -1.6453077 | 0.003299613 |
| *ISG15* | -1.6554255 | 0.002552402 |
| *CCL4L1* | -1.6829141 | 0.030765338 |
| *CD79A* | -1.7055949 | 0.003934981 |
| *HELZ2* | -1.771763 | 0.001550459 |
| *NHLH1* | -1.7918022 | 0.047734657 |
| *CMPK2* | -1.8613222 | 0.006869981 |
| *MOXD1* | -1.8928461 | 0.019198944 |
| *S100B* | -2.0415679 | 0.029057755 |
| *CCL3L1* | -2.0616965 | 0.035218182 |
| *IFIT3* | -2.2076466 | 0.000917284 |
| *CLEC17A* | -2.2947181 | 0.029525951 |
| *NTN1* | -2.5810615 | 0.010594093 |
| *IFIT2* | -2.7374365 | 1.07915E-07 |
| *PLEKHG6* | -2.8344978 | 0.000265876 |
| *IFIT1* | -2.9297855 | 0.005441837 |

### SFig4. Rab5 maintains surface CAR expression by enhancing CAR endocytic recycling

A

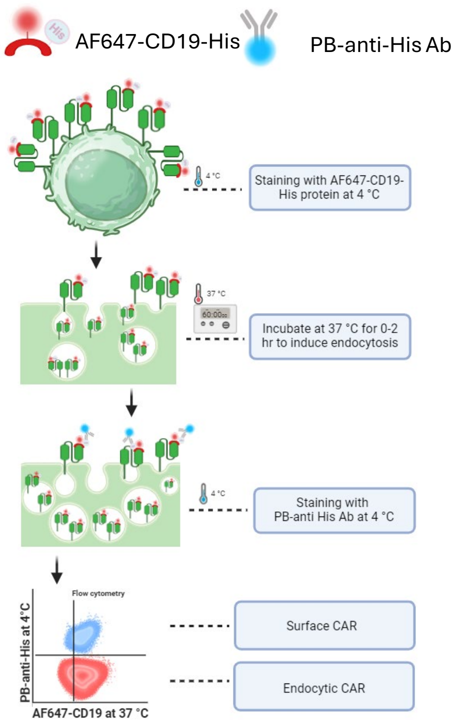


B

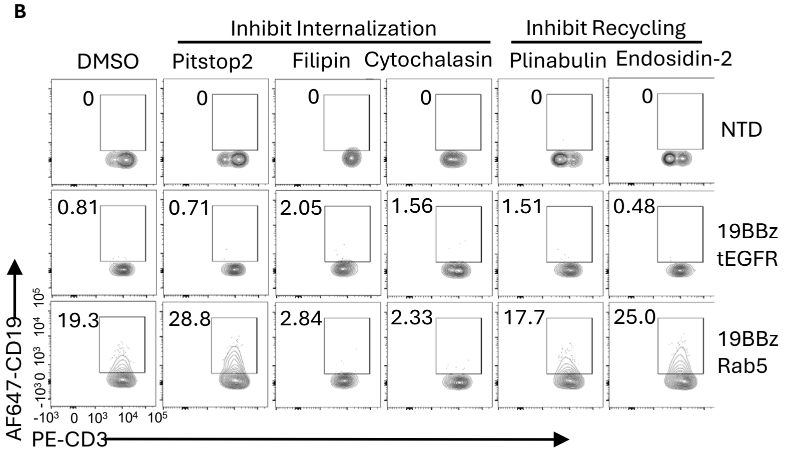


C

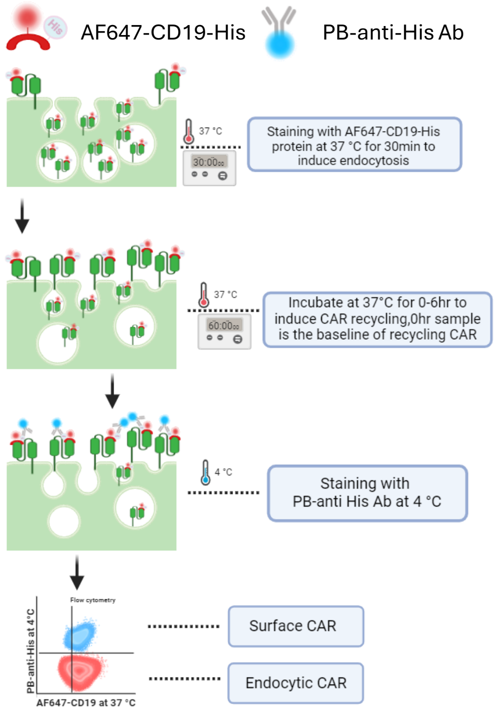

### SFig5. Rab5 CARTs exhibit superior in vivo antitumor efficacy

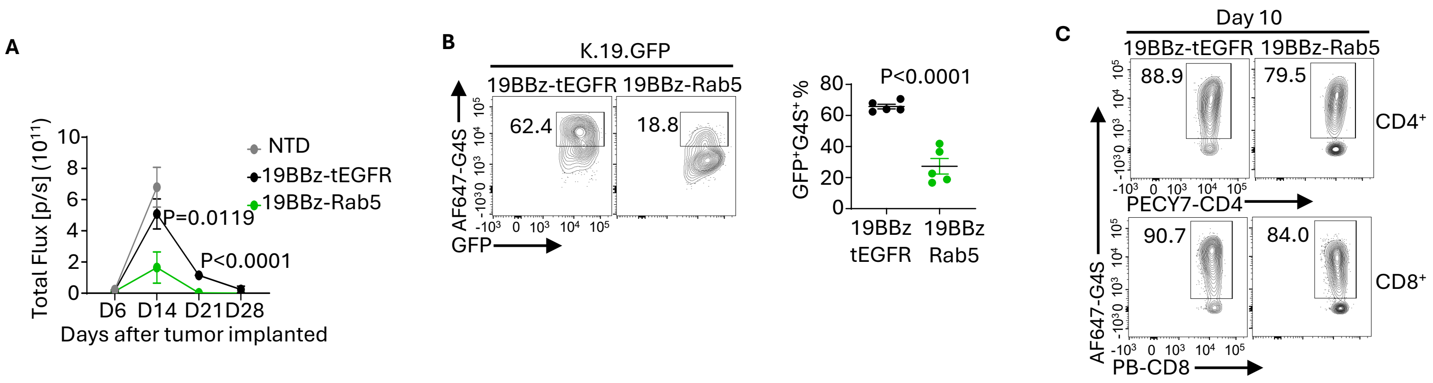

### SFig6. Rab5 expression augments the ability of CAR T cells to control solid tumors

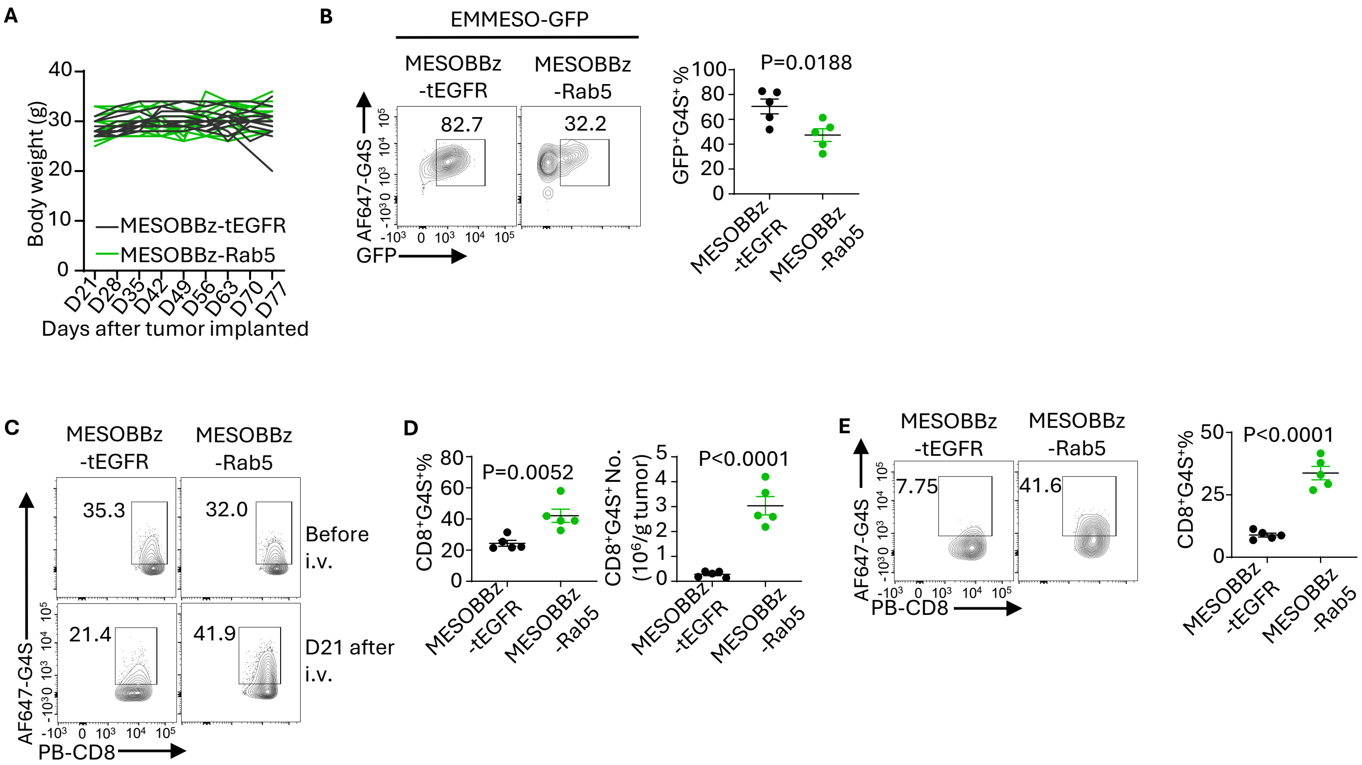

**Figure S1. CARTs continuously exposed to antigen lose CAR expression is mediated by tumor cells, related to Figure 1.**

(A). Schematic of continuous antigen exposure (CAE) assay. Primary human T cells were activated and transduced with 19BBz-CAR. After 9 days of culture, CAR expression was measured by staining with a labelled CD19 protein. At this point, $1x10^6$ CAR positive T cells were co-cultured with $2x10^6$ K.19.GFP cells. Every 2 days, additional $2x10^6$ K.19.GFP were added to the co-culture assay. Surface CAR expression was measured every day following with CD19-labeled protein staining, tumor cells killing was monitored with GFP detecting by flow cytometry.

(B). Surface CD19-CAR expression was detected using anti-G4S linker antibody after 24 hours co-culture.

(C). Summary data from three experiments showing CAR expression and tumor killing in both $CD4^+$ CARTs and $CD8^+$ CARTs throughout the CAE assay.

(D-E). Surface CAR expression downregulation is a general process. $1x10^6$ CAR positive 1928z-CARTs from day 9 were co-cultured with $2x10^6$ with K.19.GFP for CAE assay (D), or $1x10^6$ CAR positive MESOBBz-CARTs from day 10 were co-cultured with $3x10^6$ with EMMESO-GFP cells for CAE assay (E). Surface CAR expression staining with CD19-labled protein (D), or MSLN-labeled protein (E) was measured by flow cytometry.

(F-G). Long term challenge mediates surface CAR loss is tumor cells dependent. $1x10^6$ CAR positive 19BBz-CARTs were co-cultured with K.19.GFP (F), or $1x10^6$ CAR positive MESOBBz-CARTs were co-cultured with EMMESO-GFP cells (G) as indicated ratio for 24 hours, surface CAR was measured with indicated labeled protein staining by flow cytometry.

(H). Surface CAR loss directly requires interaction between CARTs and tumor cells. Same number of resting stages 19BBz-CARTs from day 11 were mixed with K.19.GFP (directly) or separately co-cultured in the transwell with K.19.GFP (indirectly) for 24 hours, surface CAR expression was examined by flow cytometry.

(I). Functional test of 19BBz-mCherry CARTs. mCherry was fused after CD3ζ in the 19BBz-CAR vector, the tumor killing function, mCherry-CAR expression and surface CAR expression was detected to confirm its functional by flow cytometry.

(J). TME imaging of APEX2-CAR (black) was shed from CARTs after tumor cells challenge. $1x10^6$ 19BBz-APEX2 CARTs were co-cultured with $2x10^6$ K.19.GFP for 24hours. T cells were isolated, followed by DAB staining for TEM.

Experiments shown in (B), and (D) to (I) are representative of three independent experiments, in (J) reproducible in two independent experiments with different donors.

**Figure S2. Rab5 enhances surface CAR persistence and CART function, related to Figure 2.**

(A-B). Flow cytometry (A) and summary data (B) of CAR surface expression is measured in $CD8^+$ CARTs in CAE assay.

(C). Schematic of CAE in vitro assay with MESO-CARTs and EMMESO-GFP cells.

Experiments shown in (B, C) is representative of three independent experiments.

**Figure S3. Rab5 regulates surface CAR expression without altering CAR stability, related to Figure 3.**

(A-F). Rab5 has no effect on CAR molecular stability. CAR stability was determined in the presence of indicated inhibitors by showing relative CAR expression with 19BBz-tEGFR or 19BBz-Rab5 CARTs that were cultured for 11 days (A-C), or the indicated CARTs were co-cultured with K.19.GFP for 48 hours after the second time of re-challenge in CAE assay (D-F).

(G-H). RNA-seq analysis was performed to assess differentially expressed genes (DEGs) between CARTs transduced with either 19BBz-tEGFR or 19BBz-Rab5 CAR. Shown are volcano plots displaying gene expression changes in 19BBz-tEGFR CARTs (control) vs 19BBz-Rab5 CARTs (test). Genes were color-coded as follows: no significant changes in expression (gray), upregulated genes with >3-fold change in expression with an adjusted p-value < 0.05 (green), downregulated genes with >3-fold change in expression with an adjusted p-value < 0.05 (black). Lists of up- and down-regulated genes are also provided. (G) Presents differences between 19BBz-tEGFR and 19BBz-Rab5 CARTs genes at baseline (after 9 days of in vitro expansion).

(H) Presents differences observed following three rounds of 19BBz-tEGFR CARTs or 19BBz-Rab5 CARTs co-cultured with K.19.GFP target cells. For both (G) and (H), N = 4; two-sided DESeq2 analysis.

In (A) to (F), reproducible in three independent experiments with different donors.

**Figure S4. Rab5 enhances CAR endocytic recycling, related to Figure 4.**

(A). Schematic of endocytic rate assay. Surface CAR was initially stained with AF647-CD19-His tag protein at 4 °C for 30min, followed by incubation of CARTs at 37 °C for indicated period to induce endocytosis. Then the surface remaining CAR was stained with PB-anti-his antibody at 4°C for 20mins, defined as AF647 and PB double positive population. The endocytic CAR was measured as the AF647 single positive population.

(B). Rab5 maintains CAR expression dependent on CIE pathway. $1x10^6$ 19BBz-tEGFR or 19BBz-Rab5 CARTs were incubated with $1x10^6$ K.19.GFP cells for 24 hours, followed by treatment with the indicated inhibitors for 30min at 37 °C. After washing, additional $1x10^6$ K.19.GFP cells were added to the co-culture for more 24 hours. Then the surface CAR was stained with CD19-labled protein and tested by flow cytometry.

(C). Schematic of recycling rate assay. Surface CAR was stimulated and stained with AF647-CD19-His tag protein at 37 °C for 30min to allow CAR endocytosis. Partial of the samples were fixed and will be used as baseline of surface and endocytic CAR expression. Partial of the samples were incubated at 37 °C for the indicated period to induce CAR recycling to the surface. At the end of recycling assay, all the samples were sainted with PB-anti-His antibody at 4°C for 20mins. The recycled CAR was identified as AF647 and PB double positive population.

Data shown in (B), reproducible in three independent experiments with different donors.

**Figure S5. Rab5 CARTs exhibit enhanced in vivo antitumor efficacy, related to Figure 6.**

(A). Tumor burden was quantified as the total photon count of each mouse, related to Figure 6B.

(B) Tumor-captured CAR was detected using intracellular staining with an anti-G4S linker antibody, following the separation of tumor cells from tumor tissues.

(C). Surface CD4 and CD8 CAR expression from 19BBz-tEGFR or 19BBz-Rab5 CARTs before infusion was tested by flow cytometry.

Data shown in (A), reproducible in three independent experiments with different donors. In (B, C), reproducible in two independent experiments with different donors. N=5 per group. Error bar show mean± SEM. P value was determined by Students t test.

**Figure S6. Rab5 expression augments CD8$^+$ CARTs to control solid tumors without side effect, related to Figure 7.**

(A). The body weight of individual tumor bearing mice was monitored per week from the day of CARTs infusion, related to Figure 7B.

(B). CARs internalized into tumor cells were determined by intracellular staining of CARs using tumor cells harvested from MESOBBz-tEGFR or MESOBBz-Rab5 CARTs treated mice.

(C-D). Surface MESO-CAR expression on CD8$^+$ CARTs before and after CARTs infusion for 21 days was measured by flow cytometry (C). Representative of percentage and absolute number of intratumor CD8$^+$CARTs (D).

(E). Surface CAR expression on CD8$^+$ CARTs with spleen samples from 21 days CARTs therapy NSG mice.

Data shown in (A), was performed in three separate experiments. In (B) to (D), reproducible

in two independent experiments with different donors, N=5 per group. Error bar show mean± SEM. P value was determined by Students t test.
